# Overcoming insecticide resistance through computational inhibitor design

**DOI:** 10.1101/161430

**Authors:** Galen J. Correy, Daniel Zaidman, Alon Harmelin, Silvia Carvalho, Peter D. Mabbitt, Viviane Calaora, Peter J. James, Andrew C. Kotze, Nir London, Colin J. Jackson

## Abstract

Insecticides allow control of agricultural pests and disease vectors and are vital for global food security and health. The evolution of resistance to insecticides, such as organophosphates (OPs), is a serious and growing concern. OP resistance often involves sequestration or hydrolysis of OPs by carboxylesterases. Inhibiting carboxylesterases could therefore restore the effectiveness of OPs for which resistance has evolved. Here, we use covalent computational design to produce nano/pico-molar boronic acid inhibitors of the carboxylesterase αE7 from the agricultural pest *Lucilia cuprina*, as well as a common Gly137Asp αE7 mutant that confers OP resistance. These inhibitors, with high selectivity against human acetylcholinesterase, and low to no toxicity in human cells and mice, act synergistically with the OPs diazinon and malathion to reduce the amount of OP required to kill *L. cuprina* by up to 16-fold, and abolish resistance. The compounds exhibit broad utility in significantly potentiating another OP, chlorpyrifos against the common pest, the peach-potato aphid *(Myzus persicae)*. These compounds represent a solution to OP resistance as well as to environmental concerns regarding overuse of OPs, allowing significant reduction of use without compromising efficacy.

## Introduction

As the world population increases, agricultural productivity is essential for sustaining food security. This has been facilitated through the use of pesticides and insecticides for protecting crops and livestock^1^. In terms of global health, insecticides are the first line of defense against many infectious diseases; they are especially important in developing countries, where insect vectors are responsible for nearly 20% of all infectious diseases^2^. Insecticide infused nets and residual spraying of dwellings are amongst the most effective means to control the spread of these diseases^3^. However, the widespread use of insecticides has effectively selected for insects that are resistant to the toxic effects^4^. Insecticide resistance is widespread and is an urgent global problem. Since the 1940s the number of insect species with reported insecticide resistance has been rapidly increasing, and recently passed 580 species^5^. Resistance renders insecticides ineffective, and leads to increased usage with significant consequences to non-target species and harm to agricultural workers^5, 6^.

Organophosphates (OPs) and carbamates are two of the most widely used classes of insecticides^7^. They inhibit acetylcholinesterase (AChE) at cholinergic neuromuscular junctions, by phosphorylating/carbamylating the active site serine nucleophile^8^. This leads to interminable nerve signal transduction and death^9^. Resistance to OPs and carbamates has been documented in many insect species^4^, with the most common mechanism of resistance involving carboxylesterases (CBEs)^10^. Resistance-associated CBEs are either overexpressed to sequester insecticides^11^, or mutated to gain a new hydrolase function^12, 13^; both mechanisms allow CBEs to intercept insecticides before they reach their target, AChE. The sheep blowfly *Lucilia cuprina* has become a model system for the study of insecticide resistance: resistance was first documented in 1966^14^, which was found to result from a Gly137Asp mutation in the gene encoding the αE7 CBE^15^. This resistance allele now dominates blowfly populations^16^, and the equivalent mutation has been observed in a range of other OP-resistant fly species^13, 17^. Recent work has shown that the wild type (WT) αE7 protein also has some protective effect against OPs through its ability to sequester the pesticides^18^.

Recent attempts to overcome insecticide resistance have focused on the development of new insecticides with novel modes of action^19, 20^. Although many of these new targets show promise, there are a finite number of biochemical targets and new targets are not immune from the problems of target site insensitivity and metabolic resistance. Synergists have been used in the past to enhance the efficacy of insecticides by inhibition of enzymes involved in insecticide detoxification; a prominent example of one of the few established insecticide synergists is piperonyl butoxide, a non-specific inhibitor of cytochrome P450s, which is used to enhance the activity of carbamates and pyrethroids^21, 22^. The idea of synergists can be taken further, to specifically target enzymes that have evolved to confer metabolic resistance, thereby restoring the efficacy of the insecticide to pre-resistance levels. Thus, CBEs such as αE7, being a relatively well-understood detoxification system, are ideal targets for the design of inhibitors to abolish insecticide resistance (**Fig. 1**).

**Figure 1.**
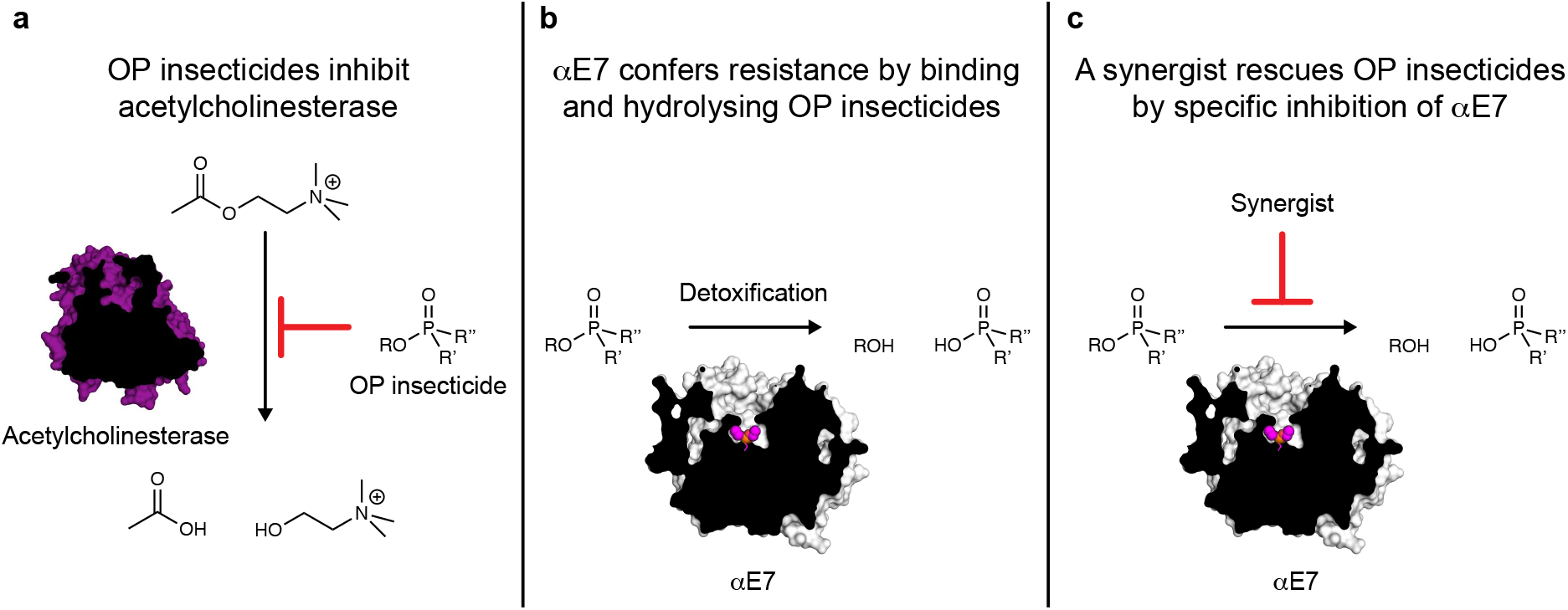
Overview of synergists for organophosphate insecticides. **(a)** Organophosphate insecticides inhibit acetylcholinesterase and prevent the hydrolysis of acetylcholine. **(b)** CBEs like αE7 rescue acetylcholinesterase activity by binding and hydrolyzing organophosphate insecticides. **(c)** An inhibitor that outcompetes organophosphates for binding to CBEs could act as a synergist to restore insecticide activity.

Here, we report the computational design of potent and selective covalent inhibitors of αE7 using DOCKovalent. DOCKovalent is a general method for screening large virtual libraries for the discovery of specific covalent inhibitors^23, 24^. An initial screen of ~23,000 boronic acids against the crystal structure of αE7^25^ identified picomolar-to-nanomolar inhibitors of WT αE7. Improving our understanding of the structure-activity relationships underlying the interaction enabled inhibitor optimization, resulting in potent inhibitors of both WT and the resistance associated Gly137Asp enzymes. Bioassays of blowfly survival confirmed that the optimized inhibitors synergized with OP insecticides and abolished resistance. Their potential for broad-spectrum use was demonstrated by the ability of the inhibitors to synergize OPs against the peach-potato aphid *Myzus persicae*. The compounds were highly selective against human AChE and did not appear toxic when administered to mice. They can overcome resistance to cheap and available insecticides, while lowering the overall amount of insecticide required by more than an order of magnitude. Such synergists could have major economic and environmental benefits and the general approach demonstrated in this work should be applicable to additional CBEs as a route to fight insecticide resistance.

## Results

### Virtual screen of boronic acids against *Lc*αE7

*Lc*αE7 catalyzes the hydrolysis of fatty acid substrates via the canonical serine hydrolase mechanism^18, 25^. Boronic acids are known to form reversible covalent adducts with the catalytic serine of serine hydrolases, which mimic the geometry of the transition state for carboxylester hydrolysis and therefore bind with high affinity^26^. We used DOCKovalent, an algorithm for screening covalent inhibitors, to screen a library of ~23,000 boronic acids against the crystal structure of *Lc*αE7 (PDB code 4FNG). Each boronic acid was modelled as a tetrahedral species covalently attached to the catalytic serine (Ser218) (**Supplementary Fig. 1**). After applying the covalent docking protocol, the top 2% of the ranked library was manually examined, and five compounds ranked between 8 and 478 were selected for testing (**Fig. 2**).

**Figure 2.**
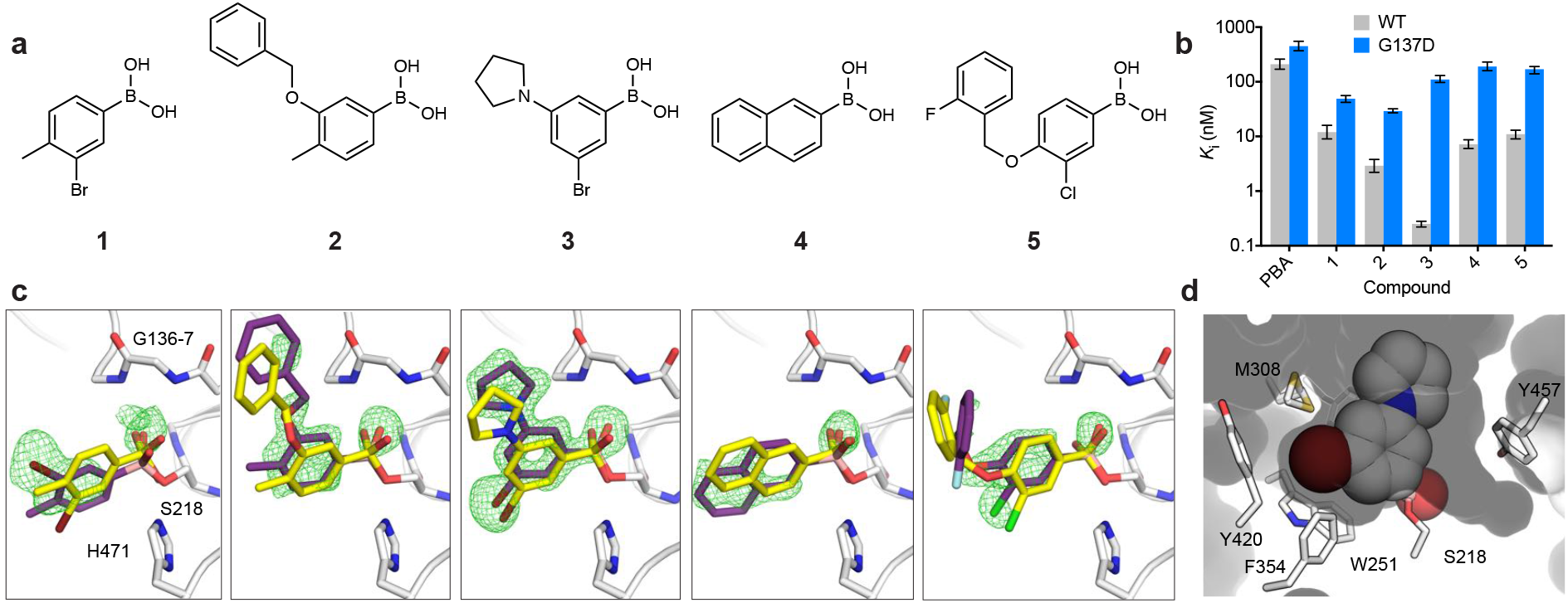
Covalent docking predicts potent inhibitors of *Lc*αE7. **(a)** Chemical structures of predicted *Lc*αE7 inhibitors. Compounds **1-5** were ranked 8^th^, 169^th^, 202^nd^, 210^th^ and 478^th^ in the DOCKovalent screen, **(b)** *In vitro K*_i_ inhibition constants for WT and Gly137Asp *Lc*αE7 with phenyl boronic acid (PBA) and compounds **1-5**. Data is presented ± 95% confidence interval for three repeat measurements of enzyme activity at each concentration of compound. **(c)** Compounds **1-5** (purple sticks) form covalent adducts with the catalytic serine of *Lc*αE7 (Ser218). The omit mF_o_-DF_c_ difference electron density is shown (green mesh contoured at 3 σ). The docking predictions (yellow sticks) are overlaid onto the corresponding co-crystal structures. Active site residues are shown as white sticks. **(d)** Surface view of *Lc*αE7 active site with compound **3** shown as spheres.

### Potent inhibitors of WT *Lc*αE7

The potency of the boronic acids was determined by enzymatic assays of recombinant *Lc*αE7 with the model carboxylester substrate 4-nitrophenol butyrate. All five boronic acids exhibited *K*_i_ values lower than 12 nM (**Fig. 2, Supplementary Fig. 2**), with the most potent compound (**3**) exhibiting a *K*_i_ value of 250 pM. While the five compounds are diverse, they all share a phenylboronic acid (PBA) sub-structure. PBA inhibits *Lc*αE7 with a *K*_i_ value approximately 2-3 orders of magnitude lower than the designed compounds (210 nM) (**Fig. 2**).

### Crystallography validates docking pose predictions

We solved the co-crystal structures of compounds 1 to 5 with *Lc*αE7 to assess the binding poses predicted by DOCKovalent (**Fig. 2, Supplementary Table 1**). Difference electron density maps show the boronic acid compounds covalently bound to the catalytic serine (**Fig. 2**). The fit of the boronic acids to the active site varies with substitution pattern; the 3,5-disubstitution of compound **3** is highly complementary while the 3,4-disubstitution of the remaining compounds results in a sub-optimal fit (**Fig. 2, Supplementary Fig. 3**). The co-crystal structure of compound **3** reveals an unexpected trigonal planar adduct rather than the typical tetrahedral adduct^27^ (**Supplementary Fig. 4**).

Comparison between the various co-crystal structures and the docked poses largely validates the DOCKovalent predictions (**Fig. 2**). Compounds **3** and **5** were accurately predicted with root mean square deviation (RMSD) of only 1.11 Å and 1.61 Å respectively. Docking predicted a flipped orientation of compound **1** with respect to the bromine substituent (RMSD 2.04 Å). For compound **2**, the prediction of the benzyloxy substituent was not accurate (RMSD 3.01 Å). Finally, the naphthalene ring of **4** was flipped relative to the docking prediction (RMSD 2.02 Å) which required a change in conformation of Met308 (**Supplementary Fig. 3**).

### Potent inhibitors of Gly137Asp *Lc*αE7

The two most common CBE-mediated insecticide resistance mechanisms involve increased protein expression, or mutation to gain new catalytic (OP-hydrolase) functions. The Gly137Asp mutation is located in the oxyanion hole and positions a new general base to catalyze dephosphorylation of the catalytic serine^15, 28^. Thus, compounds that inhibit WT *Lc*αE7 as well as this common resistance associated variant would increase the efficacy of OPs by targeting both detoxification routes. Encouraged by the activity of the boronic acids **1** to **5**, we tested the compounds against the Gly137Asp variant of *Lc*αE7 (**Fig. 2, Supplementary Fig. 5**). The most potent compound was **2**, exhibiting a *K*_i_ of 29 nM. The higher affinity of compound **2** compared to **3** (29 versus 110 nM) may be due to increased steric interaction between the rigid pyrrolidinyl substituent of compound **3** compared to the flexible benzyloxy substituent of compound **2**. Indeed, the decreased affinity of all compounds for Gly137Asp *Lc*αE7 suggests that the Asp137 side chain impedes binding. This is consistent with the higher affinity of both OP and carboxylester substrates for WT *Lc*αE7 relative to Gly137Asp *Lc*αE7^28^.

### Optimization of Gly137Asp *Lc*αE7 inhibition

To improve Gly137Asp inhibition while maintaining good WT potency, we focused on elaborating compound **3**, the most potent WT inhibitor. We purchased 12 commercially available analogues of 3-bromo phenylboronic (**Fig. 3**). These compounds were chosen because they share the 3,5-disubsitution and include small and/or flexible substituents at the 5 position. We determined *K*_i_ values for WT and Gly137Asp *Lc*αE7, and, while we did not find a more potent inhibitor of WT *Lc*αE7, six of the 12 analogs exhibited picomolar *K*_i_ values (**Fig. 3, Supplementary Fig. 2** and **5**). This establishes a stable structure-activity relationship between the 3,5-disubstituted phenylboronic acid and high affinity WT *Lc*αE7 binding. Importantly, analogs **3.9** and **3.10**, which possess the 3,5-disubstitution pattern and a benzyloxy or phenoxy substituent, exhibited a 4.4- and 6.1-fold improvement in inhibition of Gly137Asp *Lc*αE7 respectively, compared to compound **3**.

**Figure 3.**
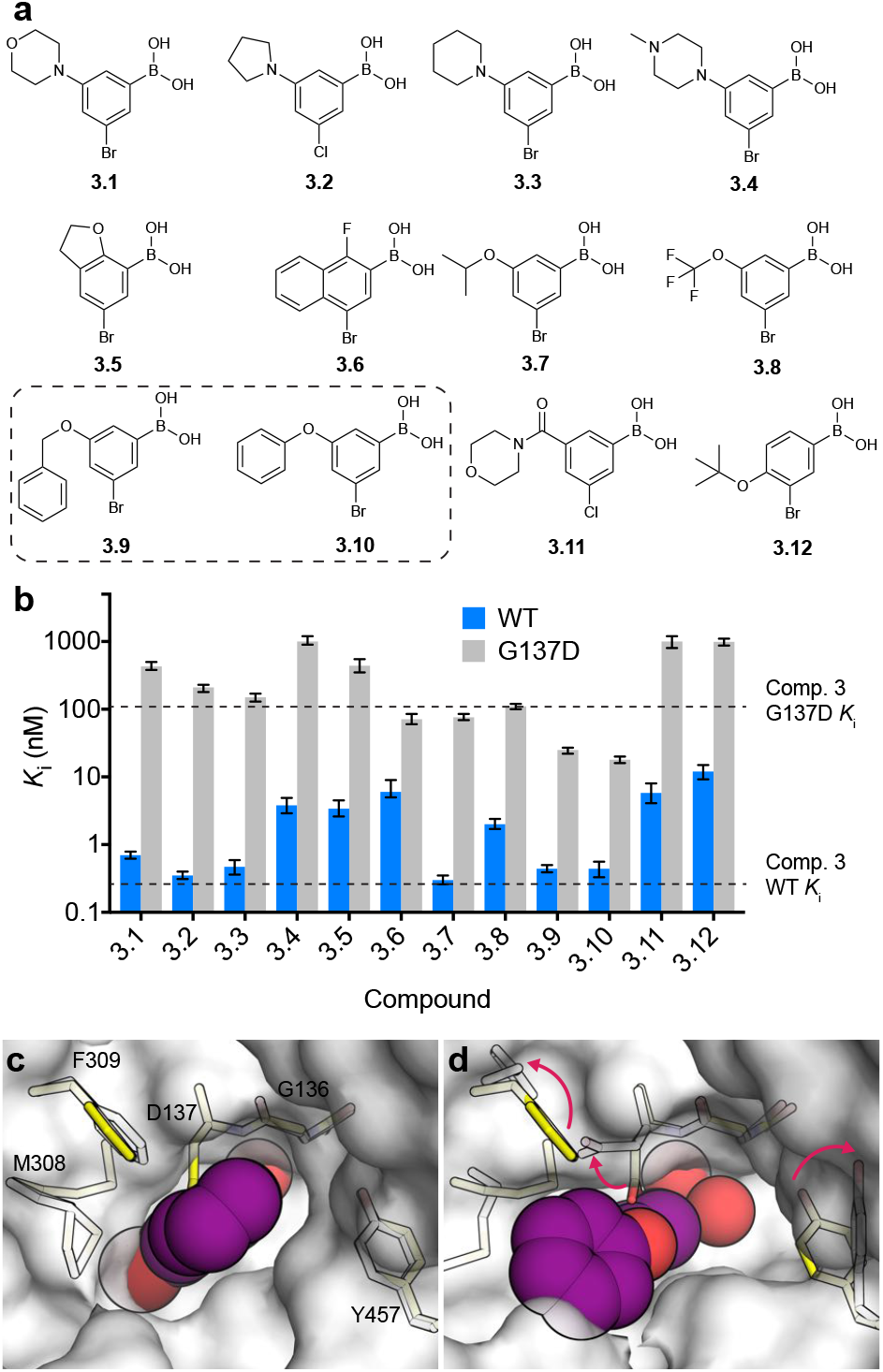
Second generation boronic acids are potent inhibitors of Gly137Asp *Lc*αE7. **(a)** Chemical structures of compound **3** analogues. **(b)** *In vitro K*_i_ inhibition constants for wild type and Gly137Asp *Lc*αE7 with compounds **3.1-3.12**. Data is presented ± 95% confidence interval for three - repeat measurements of enzyme activity at each concentration of compound. **(c)** Surface view of the co-crystal structure of compound 3 with wild type *Lc*αE7 (white surface and sticks) overlaid with the apo Gly137Asp *Lc*αE7 structure (yellow sticks). **(d)** The active site is rearranged in the co-crystal structure of compound **3.10** with Gly137Asp *Lc*αE7 (white sticks and surface). The boronic acid compounds are shown as purple spheres.

To investigate the binding of the most potent inhibitor of Gly137Asp *Lc*αE7, compound **3.10**, we determined its co-crystal structure to 1.75 Å (**Supplementary Table 1, Supplementary Fig. 6**). The orientation of **3.10** is conserved compared to **3**, with the 3-bromo substituent highly complementary to the active site, and the 5-phenoxy substituent orientated toward the active site funnel (**Supplementary Fig. 6**). The active site has undergone a reorganization, with the Asp137, Phe309 and Tyr457 side-chains adopting alternative conformations compared to the apo enzyme (**Fig. 3**). The reorganization allowed the Asp137 side-chain to avoid a steric clash with the phenoxy substituent. Surprisingly, compound **3.10** adopts a tetrahedral geometry rather than the trigonal planar geometry observed for compound **3** (**Supplementary Fig. 6**).

### Selectivity against human hydrolases

To characterize selectivity against human cholinesterases, compounds **1-5** and **3.9-3.10** were assayed against human AChE and the related blood plasma enzyme butyrylcholinesterase (BChE) (**Fig. 4** and **Supplementary Fig. 7**). The most promising compounds with respect to *Lc*αE7 inhibition (**3, 3.9** and **3.10**) were at least 20,000-fold selective for WT *Lc*αE7 over AChE. The compounds were less selective against BChE, with 1000-fold, 60-fold and 200-fold selectivity for compounds **3, 3.9** and **3.10** respectively. We also tested the compounds against human carboxylesterase 1 and 2 (CES1 and CES2) (**Fig. 4** and **Supplementary Fig. 8**). These enzymes are responsible for the majority of hydrolase activity in the liver and small intestine respectively, and their substrates include various xenobiotics and endogenous compounds^29^. Compounds **3, 3.9** and **3.10** were potent inhibitors of CES1 with *K*_i_ values in the range of 1-4 nM (**Supplementary Table 2**). Moderate selectivity was observed against CES2, with Ki values in the range of 10-1100 nM (**Fig. 4**). Notably, the *K*_i_ values for compounds 3, 3.9 and 3.10 for CES1 and CES2 is an order of magnitude less than the published IC_50_ values for widely used and irreversible OPs (*e.g*. chlorpyrifos; 0.15, 0.33 nM for hCES1 and hCES2, respectively)^30^.

**Figure 4.**
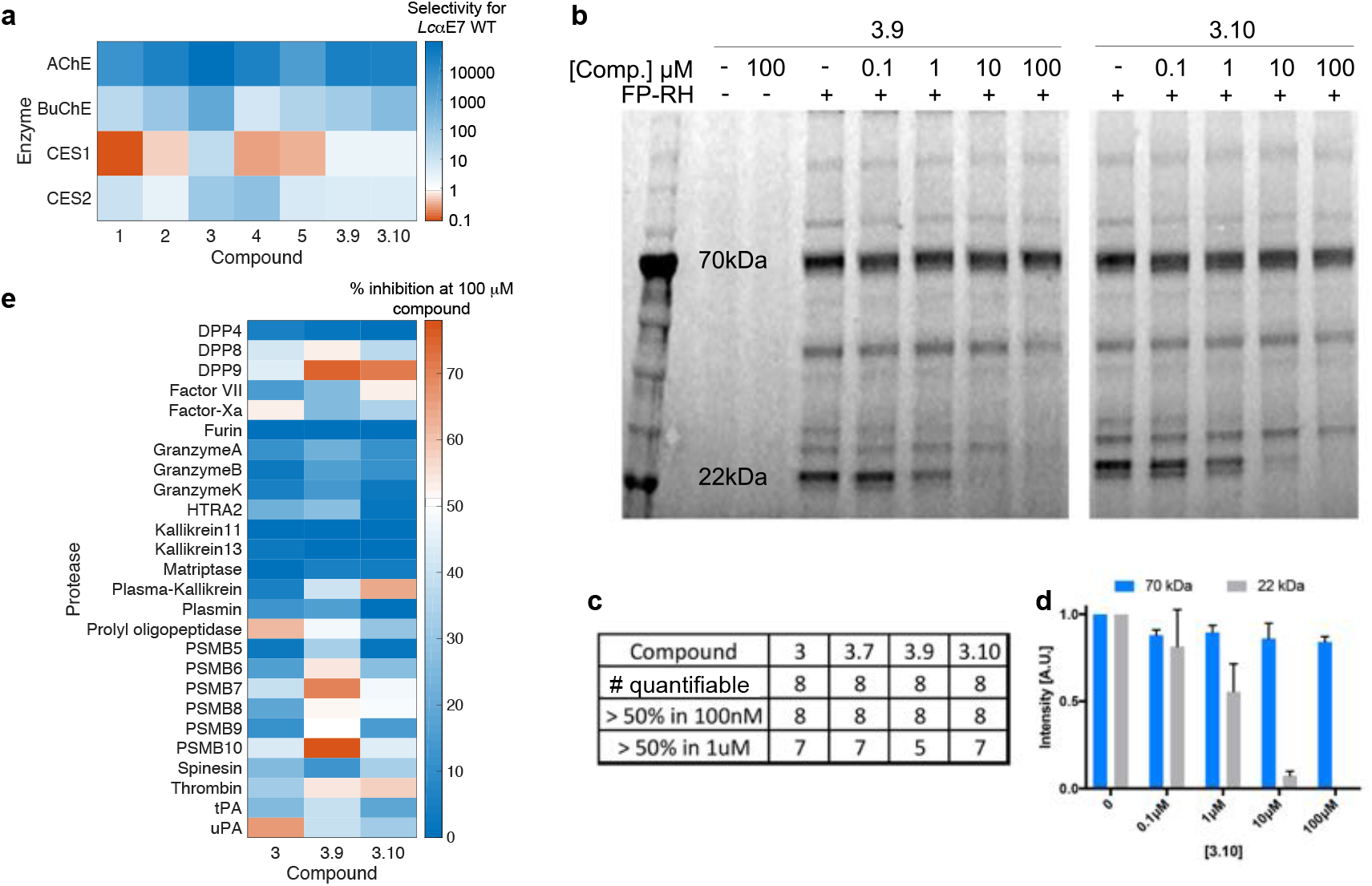
Boronic acid compounds are highly selective against human enzymes. **(a)** Heat-map showing compound selectivity against human acetylcholinesterase (AChE), butyrylcholinesterase (BChE) and carboxylesterase 1 and 2 (CES1 and CES2). Selectivity was calculated as the ratio of *K*_i_ inhibition constants between wild type *Lc*αE7 and each of the human enzymes. **(b)** In-gel fluorescence activity-based protein profiling of serine hydrolases. HEK293 cell lysates were incubated with a Fluorophosphonate-Rhodamine (FP-RH) probe, that potently and non-specifically labels serine hydrolases, with or without pre-incubation of the indicated boronic acid (10 minutes; 37°C) at increasing concentrations. **(c)** Summary of the number of quantifiable serine hydrolase bands. Only eight bands could be quantified based on these experiments. Of these, none were diminished by more than 50% with 100 nM of any of the four tested compounds. **(d)** Quantification of the two indicated bands for compound 3.10. Error bars show standard deviation from three biological repeats. **(e)** Heat-map showing inhibition of 26 human serine and threonine proteases by compound **3, 3.9** and **3.10**. Percentage inhibition was determined at a single compound concentration (100 μM) in duplicate. See **Supplementary Table 3** for assay conditions.

To explore the selectivity of the boronic acid compounds against other human serine hydrolases, we performed an in-gel activity-based protein profiling (ABPP) experiment using a fluorescent serine hydrolase probe and HEK293 cell lysate^31^ (**Fig. 4** and **Supplementary Fig. 9**). We quantified the fluorescent labelling of eight bands and found that none were diminished by more than 50% when the HEK293 cell lysate was pre-treated with 100 nM of any of the four compounds tested (**3, 3.7, 3.9** and **3.10**). We also probed the selectivity of compounds **3, 3.9** and **3.10** against a panel of 26 human serine and threonine proteases (**Fig. 4** and **Supplementary Table 3-4**). At a 100 μM concentration of the compounds, 14 out of 26 proteases retained at least 50% activity, and all proteases retained at least 20% activity. The higher concentration of compounds used in the ABPP experiment (400-fold) and protease panel (10,000-fold) relative to the *K*_i_ value of compound **3** for WT *Lc*αE7 indicates that the compounds are highly selective for *Lc*αE7 over these enzymes.

### Mammalian *ex vivo* and *in vivo* toxicity tests

Although the compounds demonstrate high selectivity against human AChE and a protease panel (**Fig. 4**), the inhibition of CES1 and CES2 indicates that off-target interactions are possible. We extended the analysis of off-target toxicity by testing the toxicity of compounds **1-5** and **3.9-3.10** against nine human cell lines (**Fig. 5** and **Supplementary Fig. 10** and **Supplementary Fig. 11**). Toxicity was assessed by the concentration of compound required to reduce cell viability to 50% (IC_50_). The compounds were generally non-toxic except against the HeLa cell line, where IC_50_ values were between 2 and 38 μM. Compound **2** was most toxic with IC_50_ values less than 50 μM for five out of nine cell lines tested.

**Figure 5.**
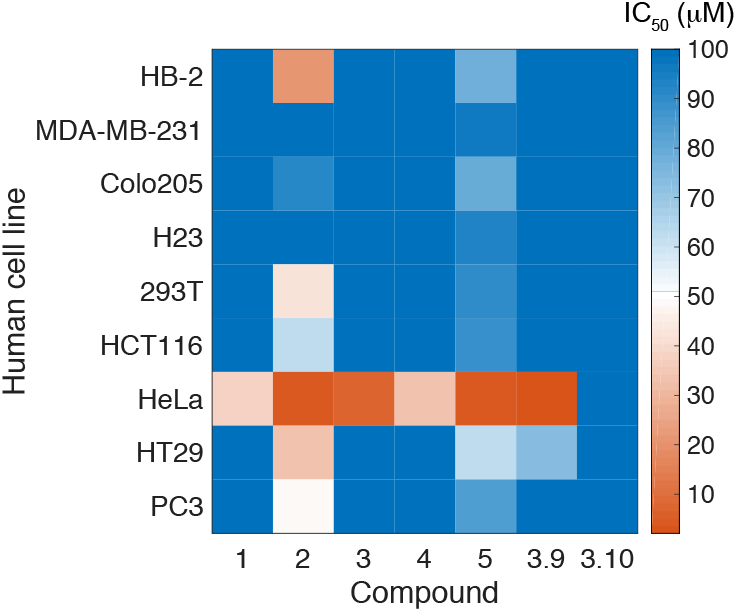
Boronic acid compounds display low toxicity against human cell lines. Heat-map showing the concentration of compounds required to kill 50% of a given cell line. The compounds were incubated with cells for 48 h and cell viability was measured using a cell luminescence assay.

We next examined the general toxicity of the boronic acid compounds using a model mammalian system of the C57BL/6 mouse. To assess mammalian toxicity, we evaluated the effects of the compounds on mice in an acute toxicity model, in which we administered the compounds at a very high dose of 300 mg/kg via oral gavage. Mice were monitored for two weeks following administration, at which point surviving mice were sacrificed and autopsied for assessment of internal organ damage. We tested compounds **3, 4, 3.7, 3.9** and **3.10** against 3-4 mice each. All mice survived and showed no clinical indication of toxicity. This demonstrates very good tolerance for the phenyl boronic acids as a general class. For context, the mouse LD50 of the OP chlorpyrifos is 62 mg/kg^32^ and the oral LD50 for Donepezil, an AChE inhibitor used clinically, is 45 mg/kg^33^. Overall, these results show that compounds from this class are relatively benign to mammals at high doses and are suitable for field use.

### *Lc*αE7 inhibitors synergistically enhance insecticides against blowfly larvae

We then investigated whether the boronic acid compounds could act as synergists to restore the effectiveness of OP insecticides. We tested the compounds against two blowfly strains; a laboratory strain (LS) which is susceptible to the OP insecticide diazinon, and the field strain “Tara”, which is resistant to diazinon^34^. Compound efficacy was determined by treating blowfly larvae with diazinon over a range of concentrations in the presence or absence of the boronic acid compounds at constant concentration, and comparing pupation^35^. Compounds **2, 3, 3.9, 3.10** and **5** were selected for testing based on high potency against WT and/or Gly137Asp *Lc*αE7, and their structural diversity. We initially tested the compounds by themselves and found that there was no significant difference between the fly pupation rates of either blowfly strain in the presence or absence of the boronic acids compounds (**Supplementary Fig. 12**). When the susceptible strain was treated with boronic acids combined with diazinon, synergism was observed for compounds **3, 3.9** and **3.10**. Compound **3.10** was the most effective, decreasing the amount of diazinon required to achieve a 50% reduction in pupation (EC_50_ value) 7.3-fold compared to a diazinon only control (**Fig. 6, Supplementary Fig. 13**). We observed similar synergism when the concentration of compound **3** was decreased from 1 mg/ml to 0.25 or 0.06 mg/ml (**Supplementary Fig. 14**).

**Figure 6.**
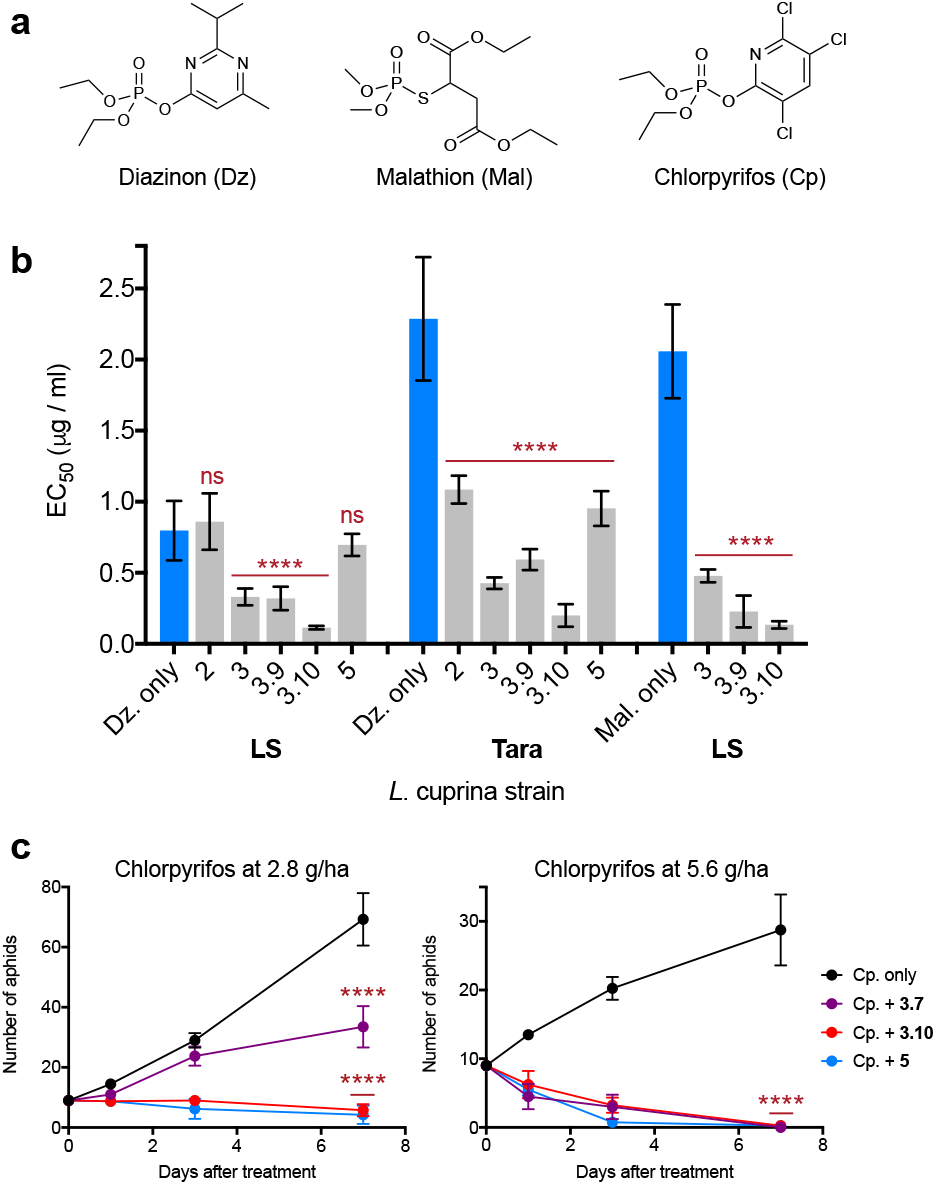
Boronic acid compounds synergize with organophosphate insecticides. **(a)** Chemical structures of the active forms of the insecticides diazinon, malathion chlorpyrifos. **(b)** Compounds **3, 3.9** and **3.10** synergized diazinon and malathion against the susceptible LS blowfly strain, while all compounds tested synergized diazinon against the resistant Tara strain. EC_50_ values were calculated from three (diazinon) or two (malathion) repeat measurements of pupation rate with 50 larvae at each diazinon/malathion concentration. EC_50_ values are presented ± 95% confidence interval with the diazinon only control compared to treatment with diazinon and the boronic acid compounds (****P<0.0001, ns = not significant; one-way ANOVA to followed by Dunnett’s multiple comparison test). **(c)** Compounds **3.7, 3.10** and **5** synergized chlorpyrifos against the peach-potato aphid. After initial infestation with 9 adult aphids, the number of alive adults and new larvae was counted after one, three and seven days. Data is mean ± SEM for four replicate experiments with the chlorpyrifos only control compared to treatment with chlorpyrifos and the boronic acid compounds (****P<0.0001; one-way ANOVA followed by Dunnett’s multiple comparison test).

Having demonstrated synergism between the boronic acids and an OP insecticide, we tested the compounds against the diazinon-resistant strain. Diazinon resistance is typically associated with the Gly137Asp mutation/resistance allele. We determined the genotype of the resistant strain, and found that, although the susceptible strain carried only WT alleles (Gly137), the resistant strain carried both WT (Gly137) and mutant (Asp137) alleles (**Supplementary Fig. 15**). This is consistent with previous reports that duplications of the chromosomal region containing αE7 have occurred, meaning that resistant strains now carry copies of both WT and Gly137Asp *Lc*αE7^36^. Resistance can be quantified by the increase in the insecticide EC_50_. The diazinon EC_50_ for the resistant strain was 2.9-fold higher compared to the susceptible strain (**Fig. 6**). This decrease in OP efficacy is within the range previously reported for eight different blowfly strains containing the Gly137Asp mutation^37^, and is sufficient to reduce OP protection in sheep from 16 weeks to 4-6 weeks^38^, and lead to failure of OPs against third instar larvae^39^. When tested against the resistant strain, all the boronic acids increased the efficacy of diazinon. The most effective compound (**3.10**) reduced the EC_50_ 12-fold. Therefore, compared to controls treated only with diazinon, compound **3.10** abolished the 2.9-fold resistance in the resistance strain, and rendered the resistant strain 4fold more sensitive to diazinon compared to the susceptible strain.

Based on the *in vitro* enzyme inhibition profile (**Fig. 2**), the higher level of diazinon synergism in the resistant strain with compound **3** versus compound **2** is surprising. However, this discrepancy is consistent with our finding that the “Tara” strain carries both WT and Gly137Asp *Lc*αE7 alleles (**Supplementary Fig. 15**); it has been shown that WT *Lc*αE7 confers substantial OP protection to the blowfly^18^. Specifically, a compound, such as **3.10** that effectively inhibits both *Lc*αE7 variants would be the best synergist against this resistant strain. The synergism exhibited by compound **3.10** is therefore a function of the optimized Gly137Asp *Lc*αE7 inhibition and retention of WT *Lc*αE7 inhibition. This highlights the importance of both sequestration *(via* WT) and catalytic detoxification*(via* Gly137Asp) by *Lc*αE7 in OP resistance.

We also tested the effects of compounds **3, 3.9** and **3.10** on the sensitivity of the laboratory strain to the OP insecticide malathion (**Fig. 6** and **Supplementary Fig. 13**). Sensitivity to malathion and diazinon is qualitatively different; WT *Lc*αE7 confers a low level of resistance to diazinon via high affinity binding and slow hydrolysis, however WT *Lc*αE7 displays significant malathion hydrolysis activity of the ester linkages in the leaving group that detoxifies the compound^13^. This difference is evident in the similar EC_50_ values for the susceptible *L. cuprina* strain treated with malathion compared to the resistant strain treated with diazinon (**Fig. 6**). Synergism with malathion was observed for all the boronic acid compounds tested. Compound **3.10** was the most effective, reducing the EC_50_ 16-fold compared to a malathion only control (**Fig. 6**).

### *Lc*αE7 inhibitors synergize with chlorpyrifos against *Myzus persicae*

To determine the potential for broad spectrum use, we tested selected boronic acids in combination with chlorpyrifos against another important agricultural pest, the peach-potato aphid *Myzus persicae*. OP resistance in *M. persicae* is mediated by two overexpressed CBE isozymes, E4 and FE4, which bind OPs with high affinity and act as insecticide “sponges”^40^. Both E4 and FE4 are related to *Lc*αE7 (50 and 53% amino acid similarity, respectively). First, we tested the toxicity of the boronic acids in the absence of OP, which confirmed the compounds have no significant toxicity by themselves (**Supplementary Fig. 16**). We then tested the efficacy of chlorpyrifos by treating adult aphids at four different concentrations, equivalent to 0.7, 1.4, 2.8 and 5.6 grams per hectare (g/ha). Significant killing only occurred at 5.6 g/ha (**Supplementary Fig. 16**). We then tested compounds 3.7, 3.10 and 5 (chosen on the basis of their structural diversity) alongside chlorpyrifos at 2.8 and 5.6 g/ha over a period of seven days (**Fig. 6**). These tests showed a significant increase in the efficacy of chlorpyrifos compared to the chlorpyrifos only controls (**Fig. 6**). While the 2.8 g/ha concentration of chlorpyrifos showed no significant killing efficacy by itself, addition of compounds **3.10** and **5** resulted in almost total extermination of the aphid population, and compound **3.7** resulted in ~50% killing. For the 5.6 g/ha concentration, the addition of all three boronic acids results in almost full efficacy of the mixture. This significant increase in chlorpyrifos efficacy, with no optimization of the compounds for *M. persicae*, establishes the broad-spectrum potential for boronic acid compounds as a new class of insecticide synergist that will allow significant reduction in the amount of OP that is required for the control of insect pests.

## Discussion

The design of potent and selective enzyme inhibitors is a central challenge in chemical biology^41^. Despite the prominence of high-throughput screening technologies, virtual compound screens provide an inexpensive and rapid alternative^42^. Here, we implemented a virtual screen of 23,000 boronic acid compounds to identify inhibitors of *Lc*αE7. The 100% hit rate from the initial screen, with all compounds exhibiting *K*_i_ values lower than 12 nM (**Fig. 2**), is surprisingly high, even when compared to previous covalent virtual screens that exhibited high hit rates against diverse enzyme targets^23^. The high hit-rate can partly be attributed to the status of boronic acid compounds as potent inhibitors of serine hydrolases^43^. Despite this general high affinity binding, the specific modification to the PBA scaffold suggested by the virtual screen afforded up to 3-orders of magnitude improvement in *Lc*αE7 inhibition. Indeed, although 74% of the screening library contained the PBA substructure, the docking still enriched it in the top hits (86% of the top 500). Therefore, although virtual screening methods will benefit from improvements to sampling and scoring, reflected by the fact that there was no correlation between docking rank and inhibitor potency for the compounds from the initial screen (**Fig. 2**), application of the DOCKovalent method allowed the rapid identification potent inhibitors of *Lc*αE7, with the initial testing of only five compounds. The initial screen also allowed us to optimize inhibitors for the Gly137Asp mutant, by combining the features of compound **3** which conferred high affinity WT binding with the features of compound **2** which were tolerated by the Gly137Asp mutant (**Fig. 2**). Further improvements to the potency and selectivity of compounds **3.9** and **3.10** may be possible by modifying the benzyloxy/phenoxy substituents using medicinal chemistry. Alternatively, 3-bromophenylboronic acid could be used as an “anchor” for the templated *in situ* synthesis of new inhibitors, as has previously been described for AChE^44^.

A major requirement for synergists to be practical is a benign safety and environmental profile. The main concern in terms of toxicity is selectivity against AChE. While the overall structures of *Lc*αE7 and human AChE are similar (PDB code 4EY4^45^, 1.05Å RMSD over 309 residues), structural differences in an active-site loop mean that the bromo-substituent of compound **3**, which is highly complementary to the *Lc*αE7 active site, is sterically occluded from the AChE active site by Phe295 (**Supplementary Fig. 17**). This results in between 20,000 to 100,000-fold selectivity for *Lc*αE7 for compounds 3 and its derivatives. In practical terms, this means that the boronic acid compounds are at least 430-fold less potent inhibitors of AChE compared to the approved drug Donepezil, an AChE inhibitor used for the treatment of Alzheimer’s disease^46^.

Having established that these compounds are unlikely to cause acute toxicity *via* inhibition of hAChE, we also tested inhibition of hBChE and hCES1/2. The structural features that confer selective binding to AChE also explain the moderate selectivity against BChE. Although AChE and BChE both possess an active-site loop that is not found in *Lc*αE7, BChE has a smaller residue (Leu286) compared to AChE (Phe295), and this difference in size reduces the steric clash at this position and therefore reduces selectivity (**Supplementary Fig. 17**). We also found low selectivity with the boronic acid compounds for *Lc*αE7 over hCES1/hCES2. The open binding pocket of CES1 (and most likely CES2, although the structure is not known) reflects its broad substrate specificity^47^ and allows the active site to accommodate the boronic acid compounds (**Supplementary Fig. 17**). Interestingly, despite the relatively open active site, CES1 possesses a similar active-site loop to AChE, and this conserved feature might allow increased selectivity to be engineered. Despite the relatively low selectivity of the boronic acid compounds for CES1, OPs are even more potent inhibitors of CES1 and are irreversible inhibitors (e.g. chlorpyrifos has an IC_50_ value of 0.15 nM with CES1^30^). Thus, the reversible inhibition of CES1/2 by the boronic acids is unlikely to cause toxicity in excess of that already observed for OPs. Furthermore, the main route of OP detoxification in humans, unlike insects, involves paraoxonases (e.g. PON1)^48, 49^ meaning that substantial synergism of OP toxicity by the boronic acids is unlikely, as PON1 is not inhibited by boronic acids^50^, and overall toxicity will be reduced because an order of magnitude less OP would be required. Follow-up studies of potential synergism could involve dosing an animal model with both OP and the boronic acids simultaneously.

To investigate whether the boronic acids might have an unexpected toxic effect, we also tested the toxicity more broadly. We showed that the compounds were selective against human serine hydrolases using in-gel ABPP (**Fig. 4**). We also demonstrated high selectivity against a panel of 26 serine and threonine proteases (**Fig. 4**) and showed that the compounds have low toxicity against nine human cell lines (**Fig. 5**). The benign nature of the boronic acids was best demonstrated by their tolerance in mice; all tested compounds were fully tolerated at a high dose of 300 mg/kg. The risk of off-target effects of the boronic acids, including the possibility that they might synergize other esterified drugs, should be considered along with the potent, non-selective and irreversible inhibition of serine hydrolases by OP insecticides. Because the boronic acid compounds would be used in combination with OPs, any toxicity associated with altered drug metabolism via esterase inhibition would be outweighed by the effect of cholinergic toxicity caused by the OP insecticides. Cholinergic toxicity is the main concern regarding OP insecticide use and the synergists described in this work could allow an order-of-magnitude reduction in OP use without compromising efficacy. Moreover, there are already examples, such as the use of piperonyl butoxide as a synergist for pyrethroids, where compounds that inhibit human metabolic enzymes (cytochrome P450s in the case of piperonyl butoxide^22^) have been safely used for decades. Given that OP insecticides are used worldwide, with an estimated 9 million kilograms applied annually in the United States alone^7^, the reduction in OP use enabled by the boronic acid synergists could have enormous health, environmental and economic benefits.

Insecticides remain the primary measure for control of agricultural pests, such as the sheep blowfly and aphids, as well as disease vectors, such as mosquitoes. The constant evolution of pesticide resistance in almost all species makes the development of new approaches to prevent or abolish resistance of great importance. While there is hope for the development of new pesticides, there are a finite number of biochemical targets and the use of synergists to knock out the resistance mechanisms and restore the effectiveness of OP insecticides is a viable alternative strategy. The compounds presented here increased the efficacy of diazinon by 12-fold against *L. cuprina* which supports the utility of targeting CBE-based resistance mechanisms. CBEs have been associated with over 50 cases of pesticide resistance over the last 50 years^10^. In addition to the high sequence conservation of metabolic insect CBEs^10^, the Gly>Asp mutation has been found in the equivalent position in two other insect pests^13, 17^, indicating that there is the potential for boronic acid based synergists to have broad spectrum activity against a range of insect species. This potential was confirmed by the ability of the boronic acid compounds to synergize with OPs against the peach-potato aphid, *M. persicae* (**Fig. 6**). An added benefit of boronic acid synergists is the potential protection from the evolution of resistance. Since boronic acids are transition state analogues for the phosphorylation of the catalytic serine by OP insecticides, mutations that hinder boronic acid binding will also likely disrupt OP sequestration and/or hydrolysis. Despite this protection, other resistance mechanisms (although far less common than CBE-mediated resistance) could still render OP insecticides ineffective, such as mutations to AChE that desensitize the enzyme to OPs^51^.

The potent and selective CBE inhibitors reported in this work represent a milestone in the use of virtual screening for inhibitor discovery in the context of combating pesticide resistance. We identified high affinity boronic acid inhibitors of a key resistance enzyme, and developed our understanding of the general structure-activity relationships that underlie the effectiveness of boronic acids with serine hydrolases, facilitating inhibitor optimization. The demonstration that the compounds effectively abolished OP insecticide resistance in *L. cuprina*, and were also effective against *M. persicae*, establishes the viability of this synergist-focused approach to combat pesticide resistance and restore the effectiveness of existing pesticide classes. The substantial increase in insecticide efficacy would allow more sustainable pesticide usage and reduce off-target environmental and health-related pesticide effects.

## Acknowledgements

We thank Paul Carr for assistance with X-ray crystallography and the Australian Synchrotron for beam time. This research was supported by the Australian Research Council (Future Fellowship to C. J. J.; FT140101059), Australian Science and Industry Endowment Fund (C. J. J. and P. D. M.; PF14-099), an Australian Postgraduate Award (G. J. C.) the Israel Science Foundation (grant No. 1097/16) to N.L. and the German-Israeli Foundation (grant No. I-2483-302.5/2017) to N.L. N.L. is the incumbent of the Alan and Laraine Fischer Career Development Chair. We thank Benjamin Cravatt for generously providing us the fluorescent serine hydrolase activity-based probe.

## Author contributions

N.L. and C.J.J. directed the project. N.L. and D.Z. performed covalent docking, computational analysis and ABPP assays. G.J.C. determined the crystal structures and performed the inhibition assays of *Lc*αE7, AChE, BChE, CES1 and CES2. P.J.J. and A.C.K. performed blowfly assays. A. H. performed mice toxicity assays. S.C. performed cellular toxicity assays. V.C. performed the peach-potato aphid assays. G.J.C., P.D.M., N.L. and C.J.J. wrote the paper. All authors contributed to the manuscript in its final form.

## Materials & Correspondence

Correspondence and requests for materials should be addressed to N.L. or C.J.J.

## Data availability

All crystal structures reported here have been deposited in the PDB under accession codes 5TYP, 5TYO, 5TYN, 5TYL, 5TYK, 5TYM and 5TYJ. PDB validation reports are all available at www.rcsb.org. All relevant data are available from the corresponding author upon request.

## Methods

### Covalent virtual screen

DOCKovalent is a covalent adaptation of DOCK3.6. Given a pre-generated set of ligand conformations and a covalent attachment point, it exhaustively samples ligand poses around the covalent bond and selects the lowest energy pose using a physics-based energy function that evaluates van der Waals and electrostatics interactions as well as penalizing for ligand desolvation. For the docking performed in this work, a boronic acids library of ~23,000 commercially available compounds was used (available for docking via http://covalent.docking.org/).

#### Receptor preparation

PDB code 4FNG was used for the docking. Ser218 was deprotonated to accommodate the covalent adduct and the Oγ partial charge was adjusted to represent a bonded form. His471 was represented in its doubly protonated form.

#### Sampling parameters

The covalent bond length was set to 1.5 ± 0.1 Å and the two newly formed bond angles to Cβ-Oγ-B=116.0 ± 5° and Oγ-B-Ligatom=109.5 ± 5°, the boron atom was replaced by a carbon, as boron is not parameterized for some of the ligand preparation tool-chain. As the covalent attachment atom is not scored during the docking, this replacement should not influence the results.

#### Candidate selection

the top 500 molecules from the ranked docking list, sorted by calculated ligand efficiency (docking score divided by number of heavy atoms) were manually inspected for exclusion, based on considerations that are orthogonal to the docking scoring function such as novelty of the compounds, diversity, commercial availability, correct representation of the molecule and internal strain (ligand internal energy is not part of the scoring function). Additionally, we selected poses in which either of the boronic acid hydroxyls is predicted to occupy the oxyanion hole. Second generation compounds were selected based on the identified 3,5 phenyl-boronic acid substitution pattern from the CombiBlocks catalog.

### Enzyme expression and purification

His_6_-tagged proteins were expressed in BL21(DE3) *E. coli* (Invitrogen) at 24°C for 18 hours. Cells were collected by centrifugation, resuspended in lysis buffer (300 mM NaCl, 10 mM imidazole, 50 mM HEPES pH 7.5) and lysed by sonication. Cell debris was pelleted by centrifugation and the soluble fraction was loaded onto a HisTrap-HP Ni-Sepharose column (GE Healthcare). Bound protein was eluted with lysis buffer supplemented with 300 mM imidazole. Fractions containing the eluted protein were concentrated with a 30 kDa molecular mass cutoff centrifuge concentrator (Amicon) and loaded onto a HiLoad 26/60 Superdex-200 size-exclusion column (GE Healthcare) pre-equilibrated with 150 mM NaCl, 20 mM HEPES pH 7.5. Eluted fractions containing the monomeric protein were pooled for enzyme inhibition assays or crystallization. Protein concentration was determined by measuring the absorbance at 280 nm with an extinction coefficient calculated using the Protparam online server^1^.

### *L. cuprina*αE7 *in vitro* inhibition assays

Inhibition of WT *Lc*αE7 and the Gly137Asp *Lc*αE7 variant was determined by a competition assay between the native-substrate analogue 4-nitrophenol butyrate (4-NPB, Sigma) and the boronic acid compounds. Initially, the Michaelis constant (*K*_M_) with 4-NPB was measured to determine an appropriate concentration of substrate for use in the competition assays. Reactions consisted of 178 μl assay buffer (100 mM NaCl, 20 mM HEPES pH 7.5), 2 μl substrate in methanol (final concentrations ranged from 1000 to 8 μM), and 20 μl enzyme (final concentrations were 2.5 nM for WT *Lc*αE7 and 4 nM for Gly137Asp *Lc*αE7). The enzymes were prepared in assay buffer supplemented with 4 mg/ml bovine serum albumin (Sigma, catalogue number A8806) to maintain stability. Enzyme velocity was determined by measuring the formation of the 4-nitrophenolate product of hydrolysis (405 nm) for four minutes at room temperature using an Epoch microplate spectrophotometer (BioTek). Initial rates were corrected for non-enzymatic hydrolysis and the Michaelis constant was determined by fitting the initial rates to the Michaelis-Menten equation using GraphPad Prism (**Supplementary Fig. 18**).

Enzyme inhibition with the boronic acid compounds was determined by assaying the initial rate of 4-NPB hydrolysis in the presence of either neat DMSO or the boronic acid compounds serially diluted 1:3 in DMSO. Reactions consisted of 178 μl assay buffer supplemented with substrate (final concentrations were 15 μM for WT *Lc*αE7 and 250 μM for Gly137Asp *Lc*αE7), 2 μl neat DMSO or 2 μl compound (final concentrations ranged from 100 μM to 60 pM) and 20 μl enzyme (final concentrations were 0.5 nM for WT *Lc*αE7 and 10 nM for Gly137Asp *Lc*αE7). Product formation was monitored for four minutes and the initial rates were corrected for non-enzymatic hydrolysis. To determine the concentration of boronic acid compounds required to inhibit 50% of esterase activity (IC50), a four-parameter sigmoidal dose-response curve was fitted to percentage inhibition using GraphPad Prism (**Supplementary Fig. 2** and **5**). *K*_i_ values were determined using the Cheng-Prusoff equation assuming competitive inhibition^2^.

### Human AChE and human BuChE *in vitro* inhibition assays

Inhibition of human AChE (Sigma, catalogue number C0663) and BuChE (Sigma, catalogue number B4186) by compounds **1-5** and **3.9-3.10** was assayed using the Ellman method^3^. Initially, the *K*_M_ was determined for AChE with acetylthiocholine and BuChE with butyrylthiocholine. Reactions consisted of 120 μl assay buffer supplemented with 5,5’-dithiobis(2-nitrobenzoic acid) (DTNB, Sigma), 40 μl assay buffer supplemented with substrate (final concentrations ranged from 1000 to 8 μM) and 40 μl enzyme (final concentrations were 0.1 nM for AChE and 0.6 nM for BuChE). The enzymes were prepared in assay buffer supplemented with 0.5 mg/ml BSA. Enzyme velocity was determined by measuring thiocholine formation (412 nm) and the Michaelis constant was determined as described previously (**Supplementary Fig. 17**). Human BuChE shows substrate activation at high concentrations of butyrylthiocholine^4^, hence the highest two substrate concentrations (1 and 0.5 mM) were excluded for fitting the *K*_M_.

Enzyme inhibition was determined in a similar manner to *Lc*αE7. Reactions consisted of 158 μl assay buffer supplemented with DTNB (final concentration of 300 μM) and substrate (final concentrations were 100 μM for both acetylthiocholine and butyrylthiocholine), 2 μl neat DMSO or 2 μl compound (final concentrations ranged from 2 mM to 400 pM) and 40 μl enzyme (final concentrations were 0.4 nM for AChE and 1.6 nM for BuChE). *K*_i_ values were determined as described previously (**Supplementary Fig. 7**).

### Human CES1 and CES2 *in vitro* inhibition assays

Inhibition of human CES1 (Sigma, catalogue number E0287) and CES2 (Sigma, catalogue number C4749) was determined for compounds **1-5** and **3.9-3.10**. The *K*_M_ was determined with the substrate analogue 4-nitrophenyl acetate (Sigma). Reactions consisted of 158 μl assay buffer, 2 μl substrate in methanol (final concentrations ranged from 1000 to 8 μM) and 40 μl enzyme (final concentrations were 3 nM for CES1 and 0.1 nM for CES2). Enzymes were prepared in assay buffer supplemented with 0.5 mg/ml BSA. Enzyme velocity was determined by measuring 4-nitrophenolate formation (405 nm) and the Michaelis constant was determined as described previously (**Supplementary Fig. 17**).

Enzyme inhibition was determined in a similar manner to *Lc*αE7. Reactions consisted of 158 μl assay buffer supplemented with substrate (final concentrations were 200 μM for CES1 and 100 μM for CES2), 2 μl neat DMSO or 2 μl compound (final concentrations ranged from 0.7 mM to100 pM) and 40 μl enzyme (final concentrations were 3 nM for CES1 and 0.4 nM for CES2). *K*_i_ values were determined as described previously (**Supplementary Fig. 8**).

### ABPP assays

#### Lysis

HEK293 cells were washed with media twice, collected, re-suspended in 1mL PBS and centrifuged (200 ref, 4°C, 5 minutes), also twice. Then, these were re-suspended in 750 μL RIPA buffer with 1:000 protease inhibitors (Pepstatin, Aprotonin, Leupeptin). Following five-minute incubation, samples were vortexed and centrifuged (16,000 g, 5 minutes). The supernatant was collected and frozen to −80°C. Lysate protein concentration was measured using a Thermo Fisher BCA kit, mixing 25 μL of 1:1, 1:10, 1:100 dilutions of the lysate in PBS/RIPA with 200 μL of BCA reagent (50:1 reagent A to reagent B). Standards were an array of dilutions of BSA also in PBS/RIPA).

#### FP-RH ABPP assay

The lysate was diluted to 1 mg/mL. Each sample was incubated with a different inhibitor concentration (0, 100 nM, 1 μM, 10 μM, 100 μM) for 10 minutes at 37°C after mixing by tapping. The samples that included the FP probe were then incubated with 1 μM of the probe for another 10 minutes at 37°C after mixing by tapping. Each sample had a total volume of 25 μL. After the second incubation, 8.33 μL of loading buffer was added to each sample (which also stops the reaction with the FP probe), vortexed and heated to 95°C for 5 minutes. Samples were loaded onto a protein gel (10 μL). Gels were imaged with green laser (532 nm) using a Typhoon fluorescence scanner. The bands were quantified using the Fiji software.

### Protease Selectivity Panel

Compounds were tested for inhibition of a panel of 26 Ser/Thr proteases at a single point concentration of 100 μM in duplicates by NanoSyn (Santa Clara, CA). Test compounds were dissolved in 100% DMSO to make 10 mM stock. Final compound concentration in assay was 100 μM. Compounds were tested in duplicate wells at single concentration and the final concentration of DMSO in all assays was kept at 1%. Five reference compounds, AEBSF, Carfilzomib, Granzyme B Inhibitor II, Dec-RVKR-CMK, and Teneligliptin hydrobromide, were tested in an identical manner with 8 concentration points at 5x dilutions. See **Supplementary Table 4** for assay conditions.

### Cellular Toxicity Assays

A seven-point, two-fold dose response series, with a 100 uM as the upper limit and a DMSO-only control point was generated using an Echo 550 liquid handler (Labcyte Inc.) in 384-well plates. Subsequently, the human cell lines were seeded (1000 cells/well) using a multi-drop Combi (Thermo Fisher Scientific) on top of the compounds. Plates were then incubated at 37°C and 5% CO_2_ for 48 hours upon which cell viability was assessed by adding CellTiter-Glo^®^ (Promega) to the reaction. The luminescence signal was measured on a Pherastar FS multi-mode plate reader (BMG Labtech).

### Mice toxicity

All animal experiments were approved by the IACUC of the Weizmann Institute of Science, Rehovot, Israel. C57BL/6 7 weeks old female mice were purchased from Envigo and allowed to acclimatize to the animal facility environment for 2 weeks before used for experimentation. Following acclimatization mice were gavaged with 300 mg/kg body weight, of the compound, dissolved in sesame oil. After the substance has been administered, food was withheld for 2 hours. Three animals were used for compounds **3.7, 3.10** and **4**, and four animals were used for compounds **3** and **3.9**. Animals were observed individually after dosing at least once during the first 30 minutes, periodically during the first 24 hours, with special attention given during the first 4 hours, and daily thereafter, for a total of 14 days. Following the 2 weeks, animals were euthanised and were subjected to histological examination, mice organs were fixed in 4% paraformaldehyde (BIO LAB, catalogue number 06450323), embedded in paraffin, sectioned (5 μm thick), and stained with hematoxylin and eosin.

### Crystallization and structure determination

Co-crystals of compounds **1-5** with the thermostable *Lc*αE7 variant^5, 6^ (*Lc*αE7-4a) (PDB code 5TYP, 5TYO, 5TYN, 5TYL and 5TLK) and compound **3.10** with Gly137Asp *Lc*αE7-4a (PDB code 5TYJ) were grown using the hanging drop vapor-diffusion method. Reservoir solutions contained 100 mM sodium acetate (pH 4.6-5.1) and 15-26% PEG 2000 monomethyl ether (MME) or PEG 550 MME. Inhibitors prepared in DMSO were incubated with protein (7 mg/ml in 75 mM NaCl and 10 mM HEPES pH 7.5) to achieve a 5:1 inhibitor-to-compound stoichiometric ratio. Hanging drops were set-up with 2 μl reservoir and 1 μl protein with crystals forming overnight at 19°C. For cryoprotection, crystals were briefly immersed in a solution containing the hanging-drop reservoir solution with the PEG concentration increased to 35%, and then vitrified at 100 K in a gaseous stream of nitrogen.

Diffraction data was collected at 100 K on either the MX1 or MX2 beam line at the Australian Synchrotron using a wavelength of 0.954 Å. Data was indexed, integrated and scaled using XDS^7^. High resolution data was excluded when the correlation coefficient between random half data sets (CC_1/2_)^8, 9^ decreased below 0.3 in the highest resolution shell. Phases were obtained by molecular replacement with the program Phaser^10^ using the coordinates of apo-LcαE7-4a (PDB code 5CH3^11^) as the search model. The initial model was improved by iterative model building with COOT^12^ and refinement with phenix.refine^13^. Inhibitor coordinates and restraints were generated with eLBOW^14^. Crystallographic statistics are summarized in **Supplementary Table 1**.

To determine if the mutations present in the thermostable *Lc*αE7-4a^5, 6^ influenced the orientation or mode of inhibitor binding, the surface mutations present in *Lc*αE7-4a were introduced into the WT background and the protein was tested for crystallization. Two mutations (Lys530Glu and Asp83Ala) were sufficient to allow crystallization in the same conditions as described previously (PDB code 5TYM) (**Supplementary Fig. 19**).

### *Lucilia cuprina* bioassays

Two strains of *L. cuprina* were used: 1) a laboratory reference drug-susceptible strain, LS, derived from collections in the Australian Capital Territory over 40 years ago, with no history of exposure to insecticides; and 2) a field-collected strain, Tara, resistant to diazinon and diflubenzuron^15^. The *Lc*αE7 gene was sequenced in each of the strains. Briefly, genomic DNA was prepared from 20 adult female flies from each strain using the DNeasy Blood and Tissue kit (Qiagen). PCR was performed with primers specific to the *Lc*αE7 gene^16^, and the product was cloned into a pGEM-T EASY vector (Promega). Eight clones of the susceptible strain and 10 clones of the resistant strain were sequenced using M13 forward and reverse primers.

The effect of compounds **2, 3, 5, 3.9** and **3.10** on the development of blowfly larvae in the presence of diazinon/malathion was assessed using a bioassay system in which larvae were allowed to develop from the first instar stage until pupation on cotton wool impregnated with diazinon/malathion over a range of concentrations, in the presence or absence of the compounds at constant concentrations^17^. Each experiment utilized 50 larvae at each diazinon/malathion concentration. Experiments were replicated three times for diazinon, and twice for malathion. The insecticidal effects were defined by measuring the pupation rate. The pupation rate dose-response data were analyzed by non-linear regression (GraphPad Prism) in order to calculate EC_50_ values (with 95% confidence intervals) representing the concentration of diazinon/malathion (alone or in combination with compounds **2, 3, 5, 3.9** or **3.10**) required to reduce the pupation rate to 50% of that measured in control assays. The effects of compounds **2, 3, 5, 3.9** and **3.10** was defined in two ways: 1) synergism ratio within each isolate = the EC_50_ for diazinon/malathion alone / EC_50_ for diazinon/malathion in combination with the compounds; and 2) resistance ratio = the EC_50_ for diazinon alone or in combination with the compounds for the Tara strain / EC_50_ for diazinon alone with the LS strain. Compounds were also tested without diazinon or malathion at 1 mg per assay.

### *Myzus persicae* bioassays

Peach-potato aphids (*Myzus persicae*) were reared on small broad beans plants *(Vicia faba*, Aquadulce variety) under controlled conditions (15°C/20°C night/day, photoperiod 14 h, 60% relative humidity). The bioassay was derived from the IRAC susceptibility test methods n°019. Briefly, seeds of broad beans were sown in small pots and cultivated under controlled conditions (15°C/20°C night/day, photoperiod 14 h) for 3 weeks. Healthy, large and flat leaves were selected from untreated plants and placed (abaxial face side up) on a Petri dish lid for treatment. Leaves were treated by spraying with a Track sprayer equipped with flat fan nozzles TeeJet XR110015VS and calibrated to deliver 200 L/ha at 400 kPa and 4 km/h. Three leaves were used for each repetition. The leaves were then air dried for 1 h at 20°C until completely dry. One leaf-disc of 2.5 cm of diameter was cut in each treated leaf and placed, abaxial face side up, in a Petri dish, on a thick layer of sterile water agar medium. Each leaf disk was then infested with 3 adult aphids with a paint brush. Petri dishes were sealed with a perforated lid with hole closed with nylon filter. One repetition comprised a Petri dish with three leaf discs and 9 aphids. Four repetitions were used for each condition. The petri dishes were incubated in the same controlled conditions as the breeding. Assessment of living aphids was made one, three and seven days after leaf infestation. Chlorpyrifos was tested at 0.7, 1.4, 2.8 and 5.6 g/ha alone or in combination with the boronic acid compounds at 0.2 mg/ml. Each boronic acid compound was also tested alone at 0.2 mg/ml with the final concentration of DMSO at 1%. Water and 1% DMSO were used as negative controls.

