## Supplementary information for "Overcoming insecticide resistance through computational inhibitor design"

### Supplementary Figures and Tables

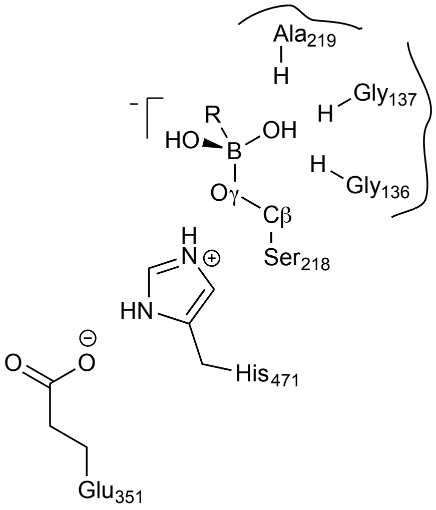

**Supplementary Figure 1. Orientation of boronic acids used for virtual screen.** The boronic acids were covalently attached to the catalytic serine (Ser218) of *Lc*αE7. The catalytic histidine (His471) was represented in its doubly protonated form. The B-Oγ bond was set to 1.5 ± 0.1 Å and the Cβ-Oγ-B bond angle was set to 116.0 ± 5° and the Oγ-B-R bond angle was set to 109.5 ± 5°. In the manual selection of compounds for testing, preference was given to compounds where either of the boronic acid hydroxyls occupied the oxyanion hole (formed by the backbone nitrogens of Gly136, Gly137 and Ala219).

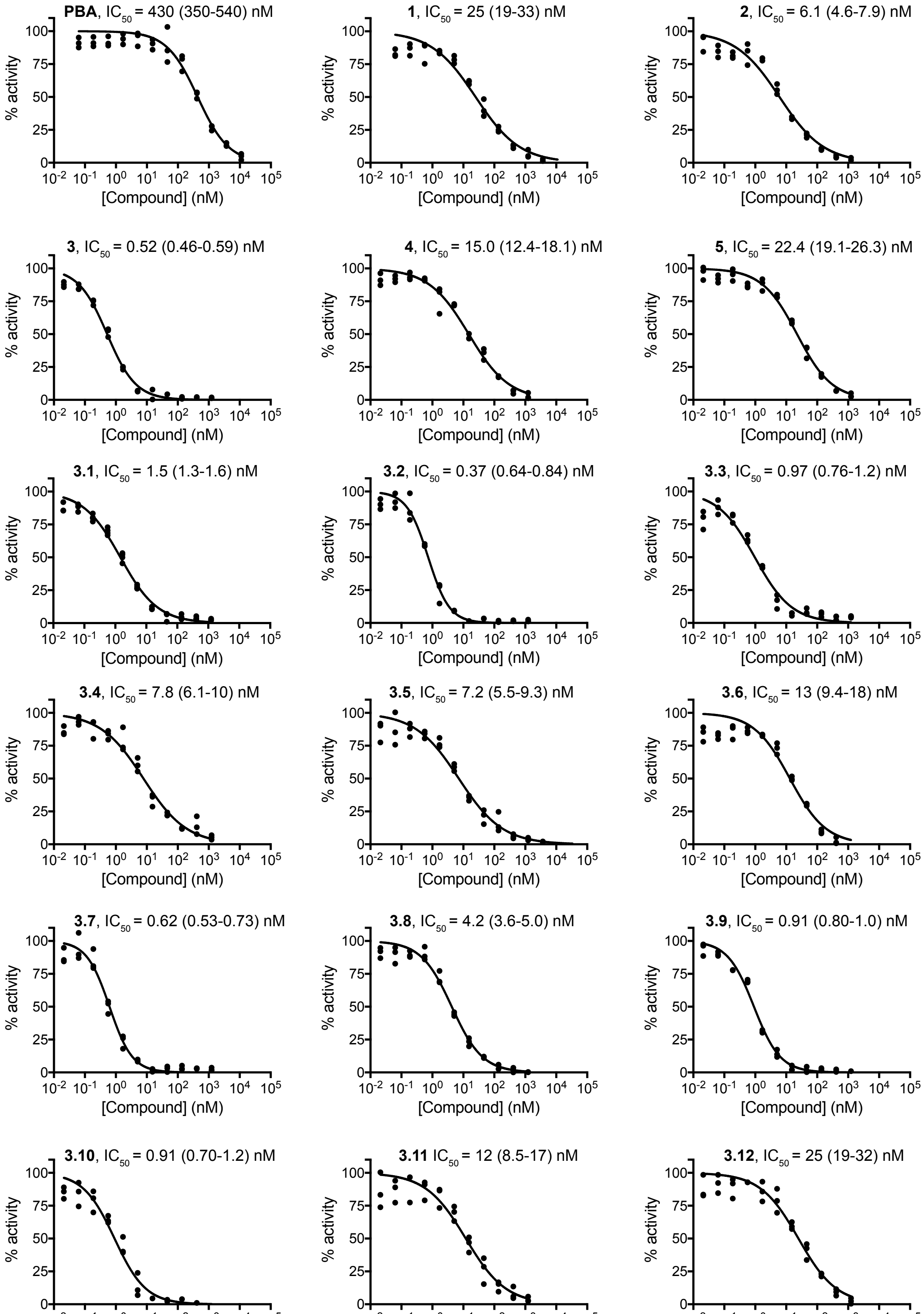

**Supplementary Figure 2. Dose-response curves for the inhibition of wild type *Lc*αE7**. Inhibition of the hydrolysis of 4-nitrophenyl was determined for phenylboronic acid (PBA) and compounds **1-5** and **3.1-3.12**. Three repeat measurements of enzyme activity were performed for each concentration of boronic acid. The concentration of boronic acid required to inhibit 50% of activity ($IC_{50}$) was determined by fitting a sigmoidal dose-response curve to plots of percentage inhibition. The curve was constrained to 0 (bottom) and 100% (top) inhibition with a variable Hill slope. $IC_{50}$ values are quoted with the 95% confidence interval.

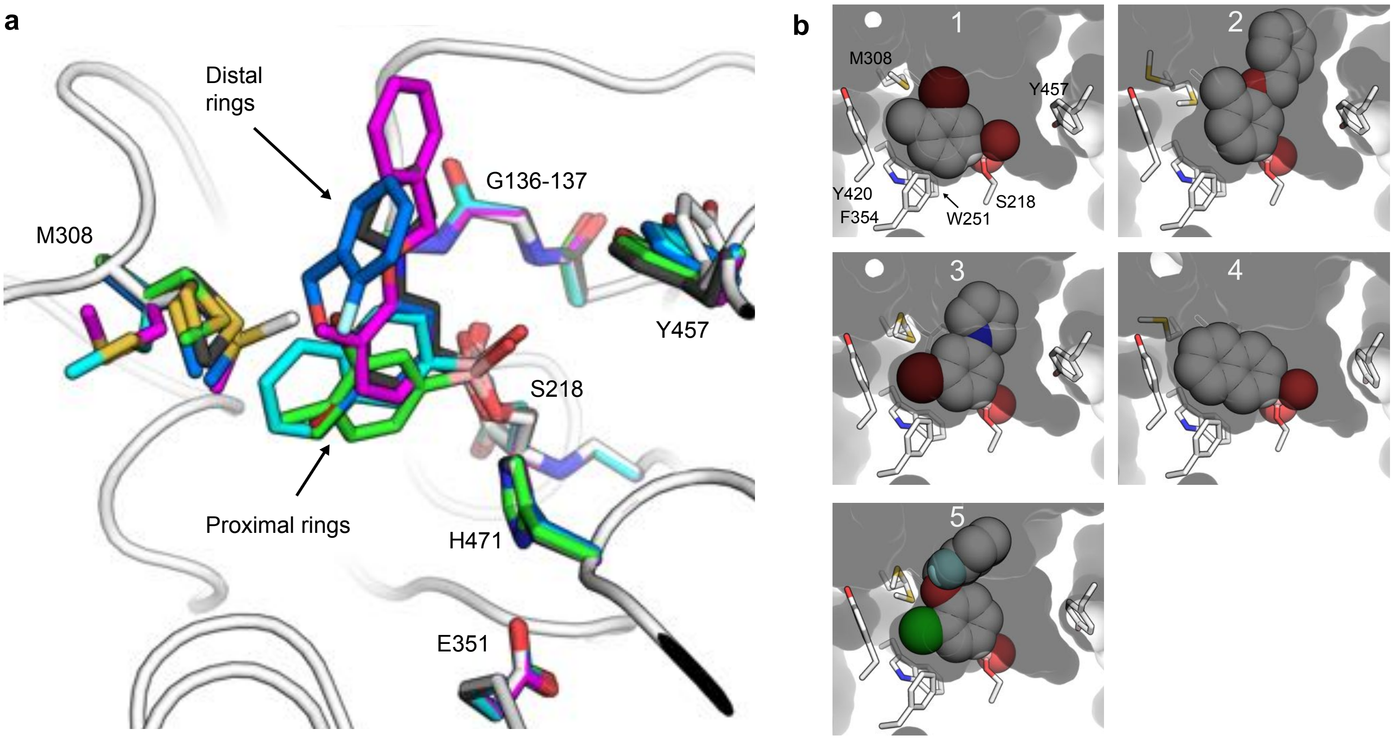

**Supplementary Figure 3. Structural features of compounds 1-5 binding to wild type *Lc*αE7. (a)** Overlay of the five wild type *Lc*αE7 co-crystal structures. The key amino acid side chains and the covalently bound boronic acids are shown as sticks coloured by inhibitor: apo = white, **1** = green, **2** = magenta, **3** = black, **4** = cyan and **5** = blue. The boronic acids occupy the larger of the two *Lc*αE7 binding pocket subsites, which accommodates a fatty acid chain of the predicted native lipid substrate[1]. The distal rings of compounds **2**, **3** and **5** are projected toward the funnel that leads to the active site. There is minor structural rearrangement in the enzyme-inhibitor complexes with the geometry of the oxyanion hole, and hydrogen bonding within the catalytic triad, remaining intact. Structural differences includes rotation of the Tyr457 side chain, which occludes the smaller subsite of the *Lc*αE7 binding pocket, and has previously been noted to occur upon OP binding[2], and changes in the position of the Met308 side-chain. **(b)** Surface representation of *Lc*αE7 binding pocket with compounds **1**-**5** shown as spheres.

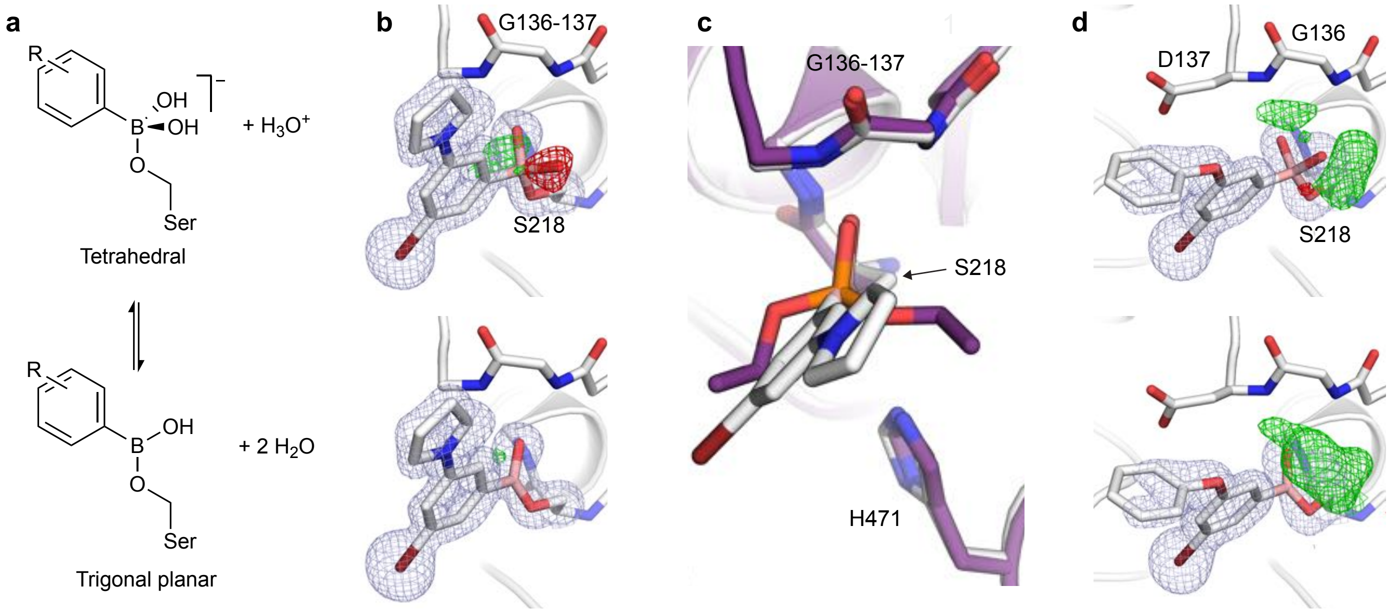

**Supplementary Figure 4. Boronic acids adopt both tetrahedral and trigonal planar geometries when coordinated to the catalytic serine of *Lc*αE7. (a)** Chemical structures of tetrahedral and trigonal planar boronic acid adducts. The tetrahedral geometry is frequently invoked to rationalize high affinity binding of boronic acid compounds to serine hydrolyase[3–8]. To our knowledge, there has only been a single report of a trigonal planar adduct[9]. **(b)** Compound **3** formed a trigonal planar adduct when bound to wild type *Lc*αE7. Geometry was assigned with reference to the positive (green mesh) and negative (red mesh) peaks in the $mF_O$-$DF_C$ difference electron density (contoured at ±3 σ) with either the tetrahedral (top) or trigonal planar (bottom) adducts modelled. The ligand and selected active site residues are shown as white sticks, with $2mF_O$-$DF_C$ electron density (blue mesh) contoured at 1 σ. Analysis of the difference density of the other compounds (not shown) indicates that compounds **2** and **5** also formed trigonal planar adducts, while compounds **1** and **4** formed tetrahedral adducts. Hydrogen bonds to the oxyanion hole were shorter for the trigonal planar adducts; the average distance was 2.7 Å for trigonal planar (compounds **2**, **3** and **5**) versus 3.0 Å for the tetrahedral (compounds **1** and **4**). **(c)** The trigonal planar adduct of compound **3** (white sticks) occupies the oxyanion hole in a similar manner compared to a bound diethyl OP (purple sticks). **(d)** Difference electron density ($mF_O$-$DF_C$ contoured at ±3 σ) suggests that compound **3.10** forms a tetrahedral adduct when bound to Gly137Asp *Lc*αE7.

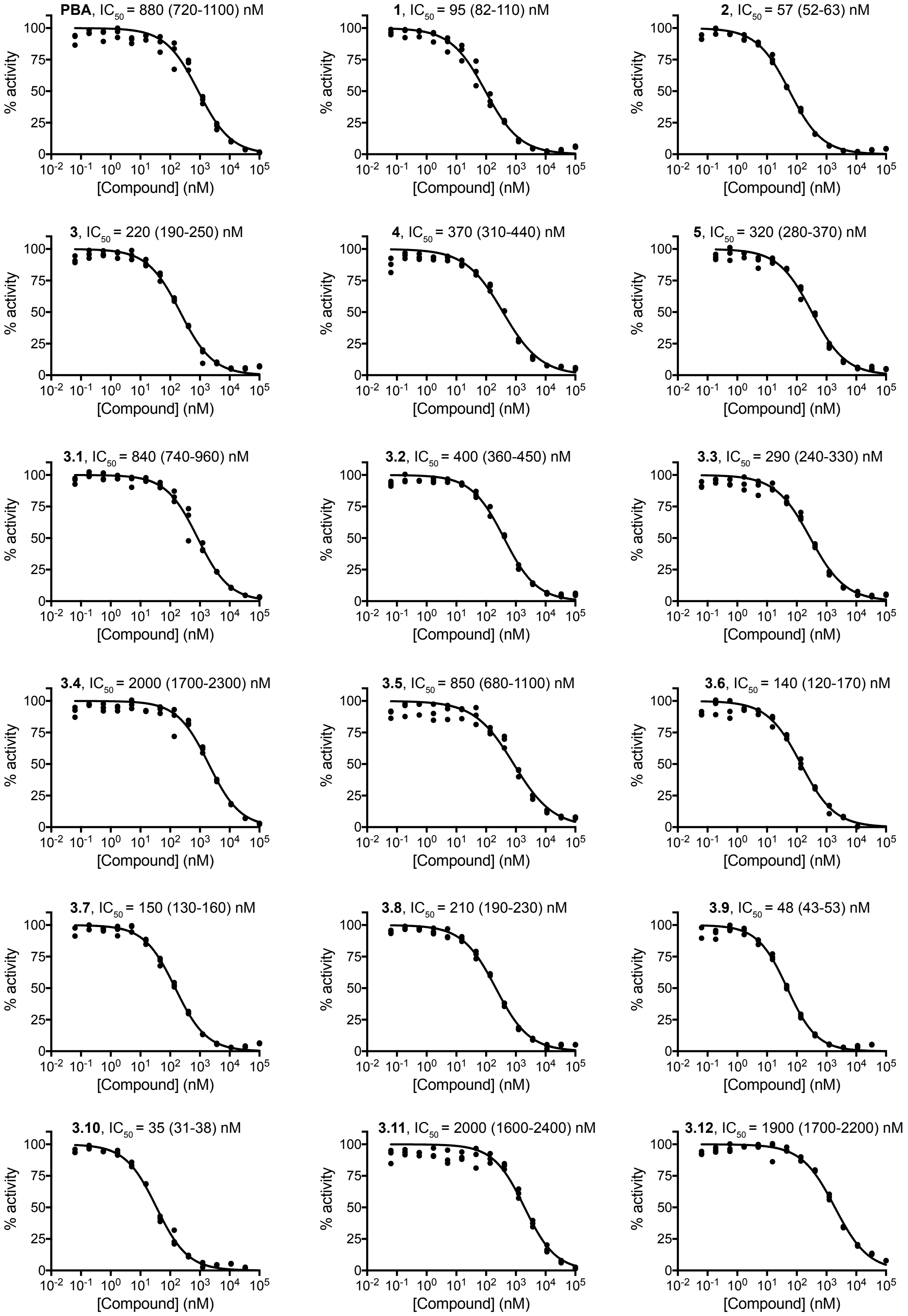

**Supplementary Figure 5. Dose-response curves for the inhibition of Gly137Asp *Lc*αE7.** Inhibition of the hydrolysis of 4-nitrophenyl was determined for phenyl boronic acid (**PBA**) and compounds **1**-**5** and **3.1**-**3.12**. Repeats and analysis was the same as Supplementary Fig 2.

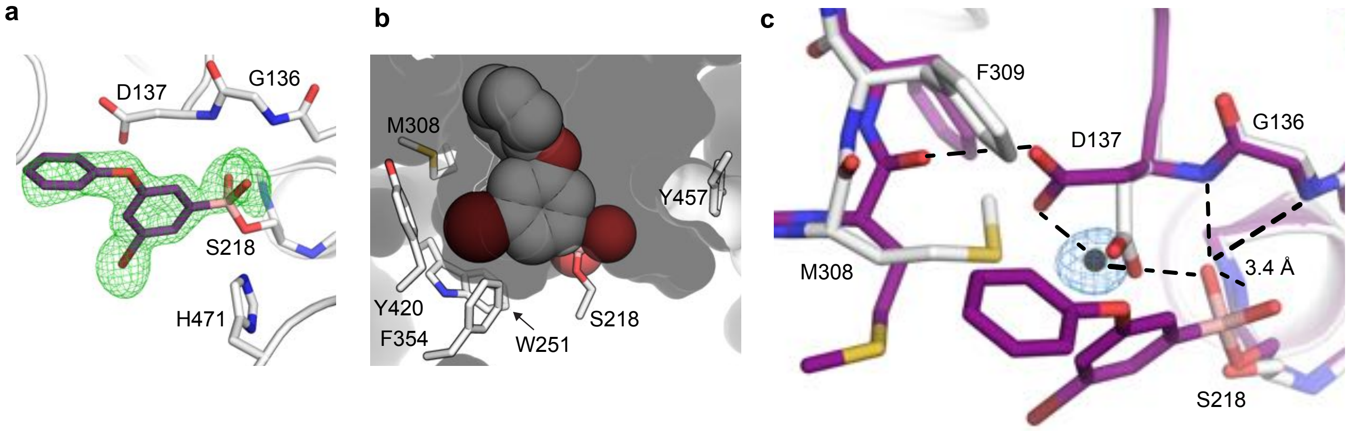

**Supplementary Figure 6. Rearrangement of the active site upon compound 3.10 binding to Gly137Asp *Lc*αE7**. **(a)** Omit density (green mesh contoured at 3 σ) showing **3.10** covalently bound to the catalytic serine (Ser218). (**b**) Compound **3.10** binding to the Gly137Asp *Lc*αE7 mutant is conserved relative to the wild type enzyme (see **Supplementary Fig. 2**). (**c**) Rearrangement of the Gly137Asp *Lc*αE7 active site upon with **3.10** binding. The co-crystal structure of compound **3.10** (purple sticks, PDB code 5TYJ) is overlaid with the apo Gly137Asp *Lc*αE7 structure (white sticks, PDB code 5C8V). To accommodate the covalently bound inhibitor, the side chains of Asp137, Met308 and Phe309 adopt new, alternative conformations. This rearrangement is consistent with molecular dynamics simulations of *Lc*αE7 which suggest that this conformational sub-state could be sampled by the free enzyme[10], and indicates that, like OPs, binding of the boronic acid **3.10** to *Lc*αE7 involves binding to minor conformational sub-states of the *Lc*αE7 active site. This rearrangement forms a new network of hydrogen bonds which connects the Met308 backbone with the bound inhibitor via the Asp137 carboxylate and a water molecule. Hydrogen bonds are shown as dashed black lines. All hydrogen bonds are 2.8 Å except for the Gly136 N to boronic acid OH which is 3.4 Å. The water molecule mediating a hydrogen bond between the Asp137 sidechain and the boronic acid is shown as a black sphere with $2mF_O\text{-}DF_C$ electron density (contoured at 1 σ).

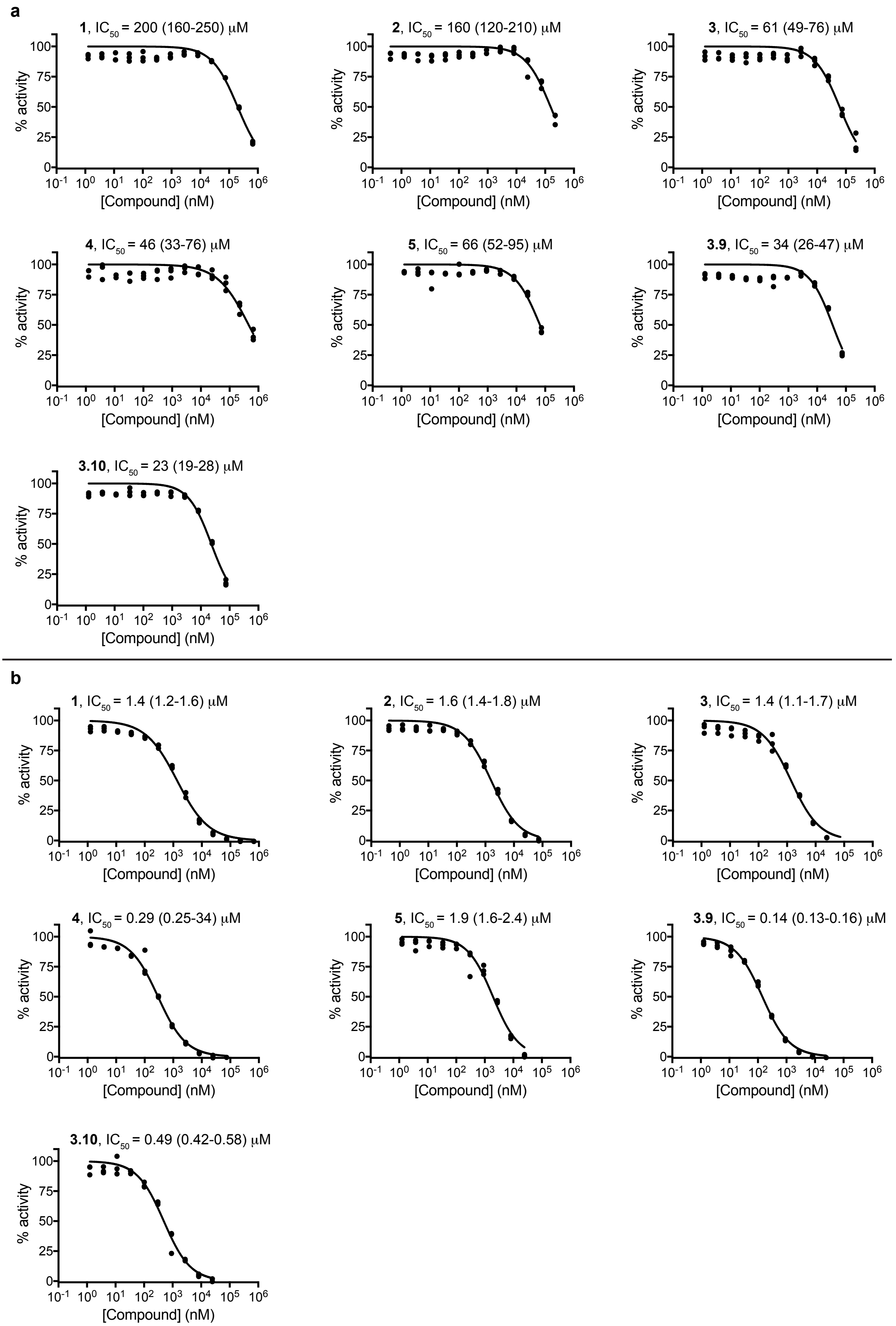

**Supplementary Figure 7. Dose-response curves for the inhibition of human acetylcholinesterase (a) and butyrylcholinesterase (b).** Inhibition of the hydrolysis of acetylthiocholine / butyrylthiocholine was determined for compounds **1-5** and **3.9-3.10**. Repeats and analysis were the same as **Supplementary Fig 2**.

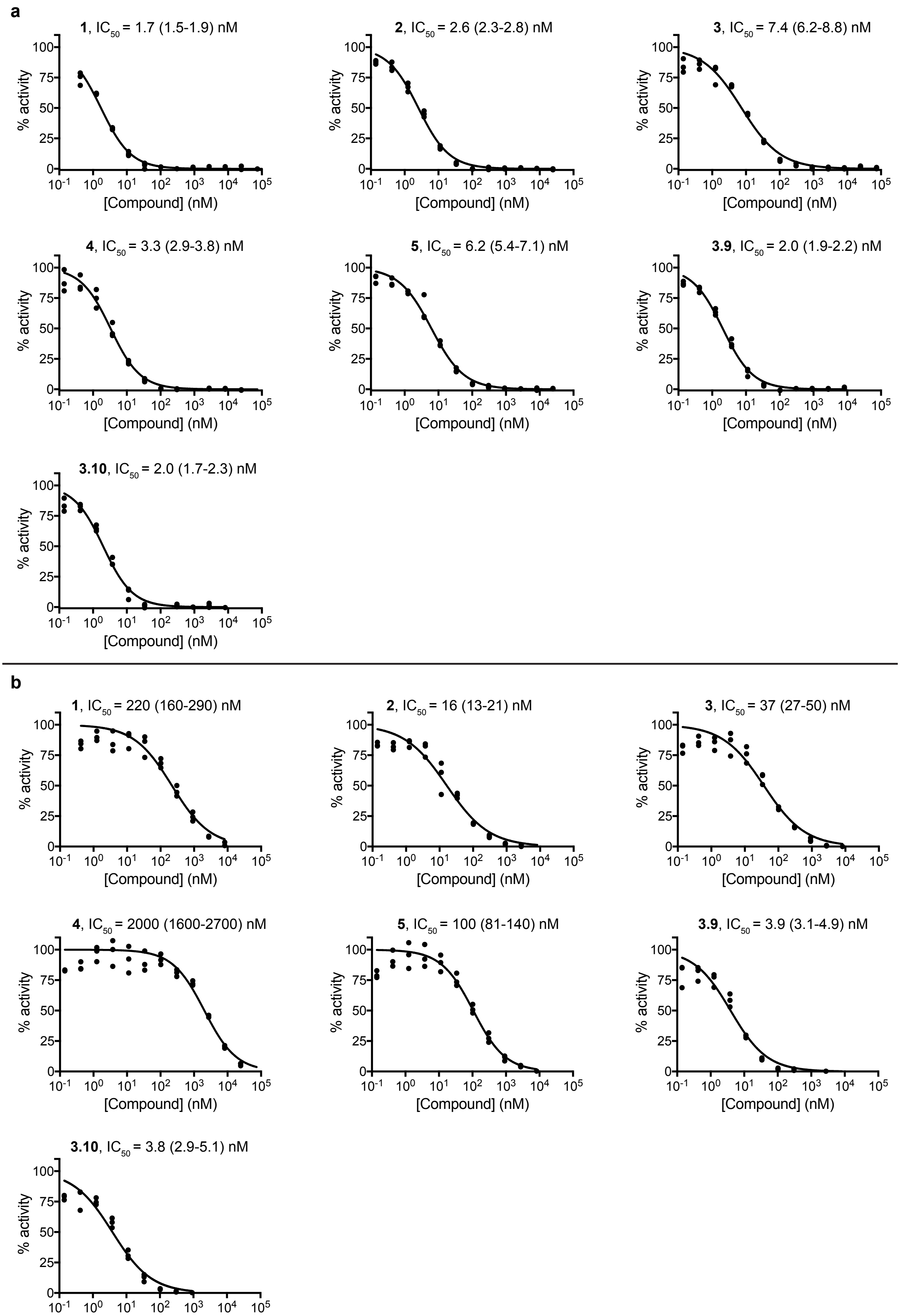

**Supplementary Figure 8. Dose-response curves for the inhibition of human carboxylesterase 1 (a) and carboxylesterase 2 (b).** Inhibition of the hydrolysis of 4-nitrophenyl acetate was determined for compounds **1-5** and **3.9-3.10**. Repeats and analysis were the same as **Supplementary Fig 2**.

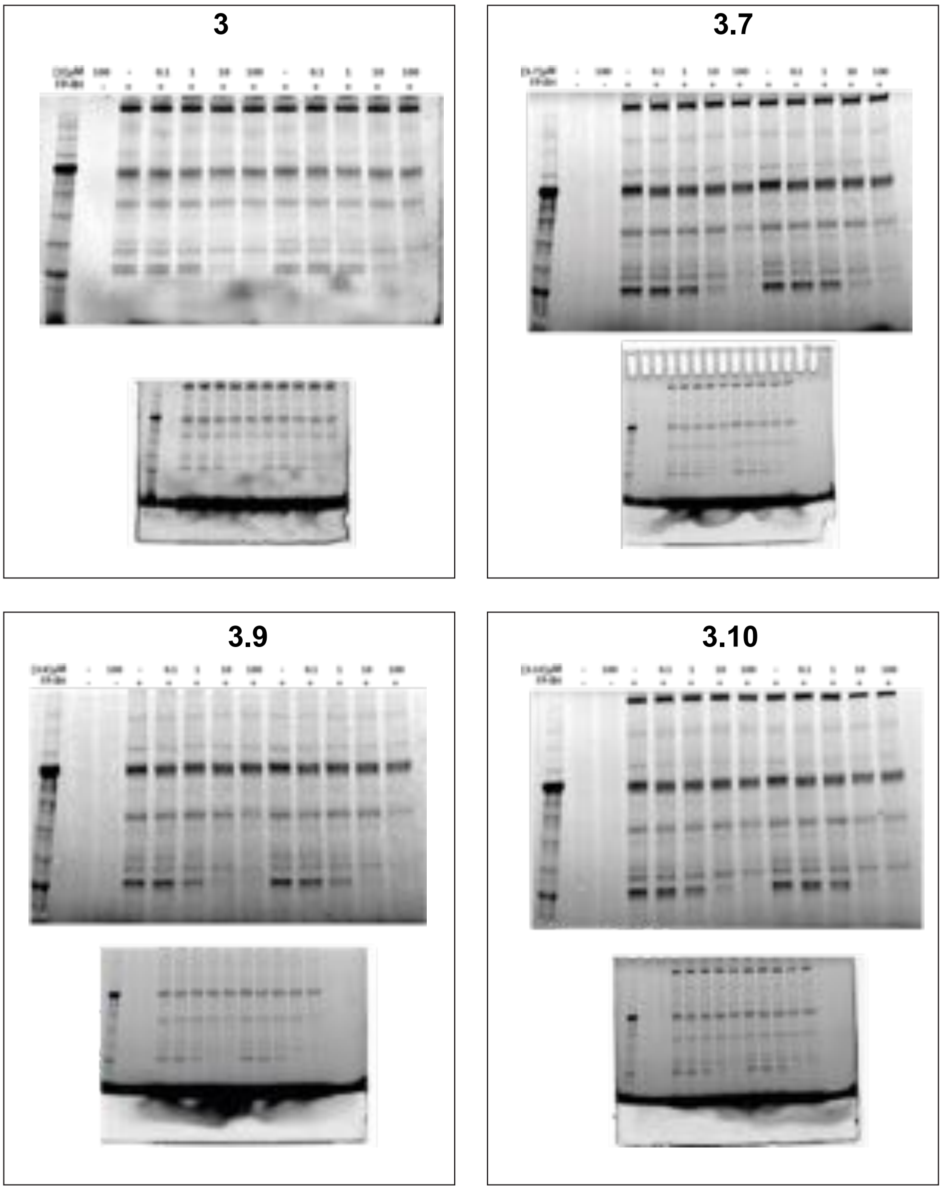

**Supplementary Figure 9. Summary of in-gel fluorescence activity-based protein profiling of serine hydrolases.** HEK293 cell lysates were incubated with a Fluorophosphonate-Rhodamine (FP-RH) probe, that potently and non-specifically labels serine hydrolases, with or without pre-incubation of the four indicated boronic acids (**3**, **3.7**, **3.9**, **3.10**; 10 minutes; 37°C) at increasing concentrations (100 nM, 1 μM, 10 μM, 100 μM). The cut and the uncut gels are supplied for those four compounds.

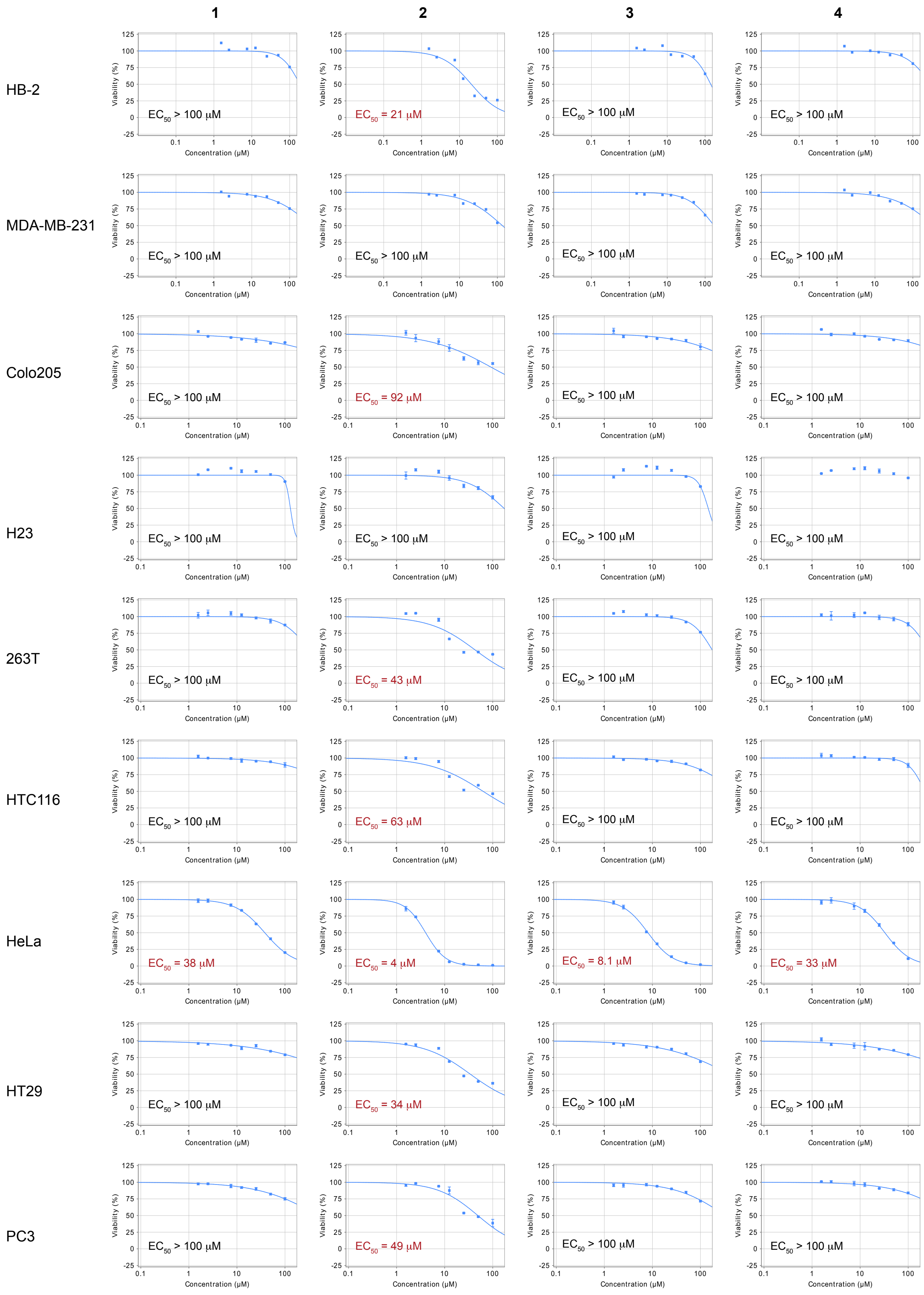

**Supplementary Figure 10. Boronic acids compounds 1-4 show little cell toxicity except against HeLa cells.** Compounds were incubated for 48 h with seven human cell lines at seven concentrations up to 100 µM. Cell viability was measured after 48 h using Cell-Titer-Glo assay. The best-fit $EC_{50}$ values are quoted on each plot for single measurements at each concentration for HB-2 and MDA-MB-231 cell lines, and duplicate measurements for the remaining seven cell lines. The cell toxicity for compounds **5**, **3.9** and **3.10** is shown in **Supplementary Fig. 16**.

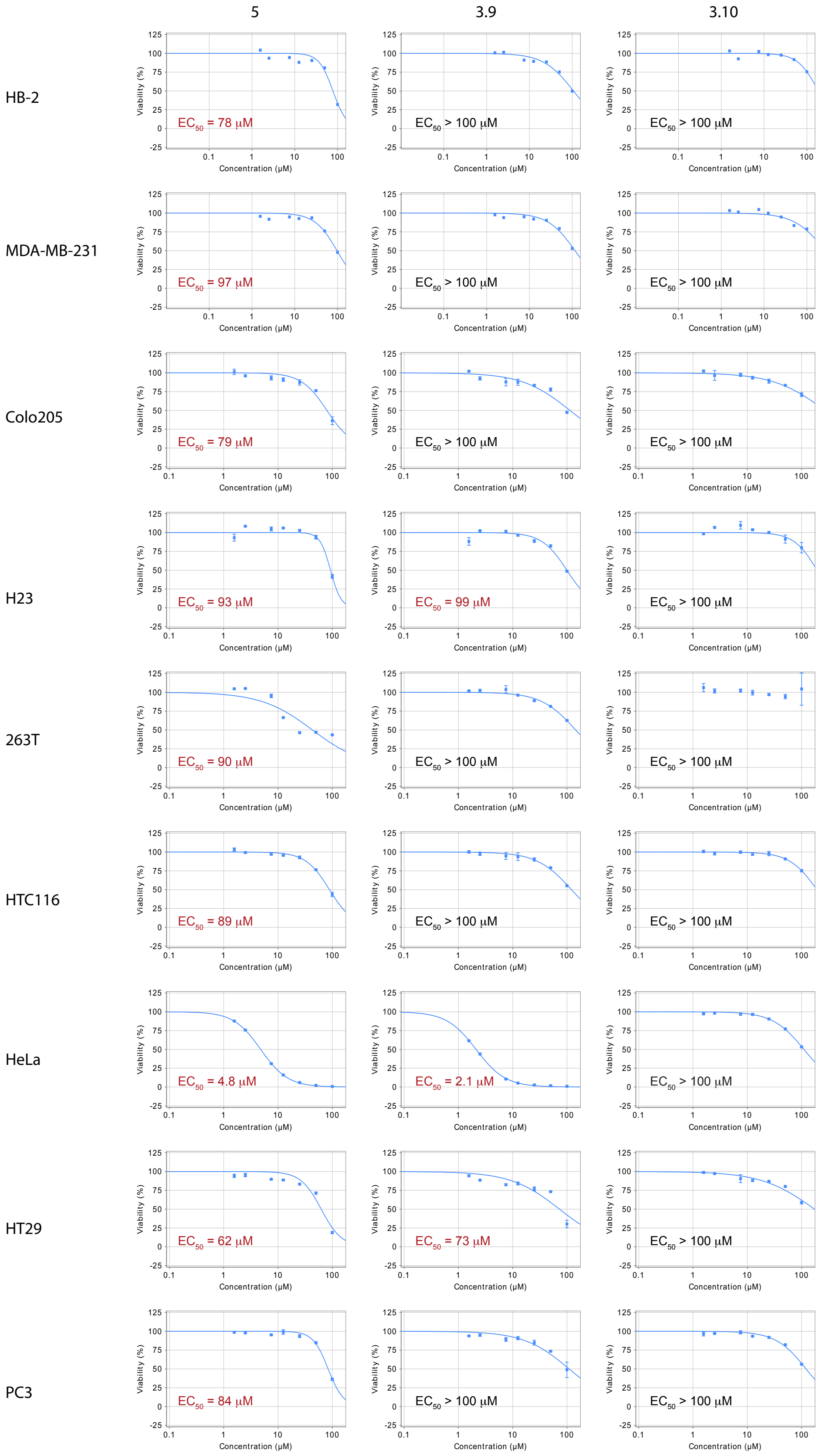

**Supplementary Figure 11. Compounds 5, 3.9 and 3.10 show little cell toxicity except against HeLa cells.** Compounds were incubated for 48 h with seven human cell lines at 7 concentrations up to 100 µM. Cell viability was measured after 48 h using Cell-Titer-Glo assay. The best-fit $EC_{50}$ values are quoted on each plot for single measurements at each concentration for HB-2 and MDA-MB-231 cell lines, and duplicate measurements for the remaining seven cell lines.

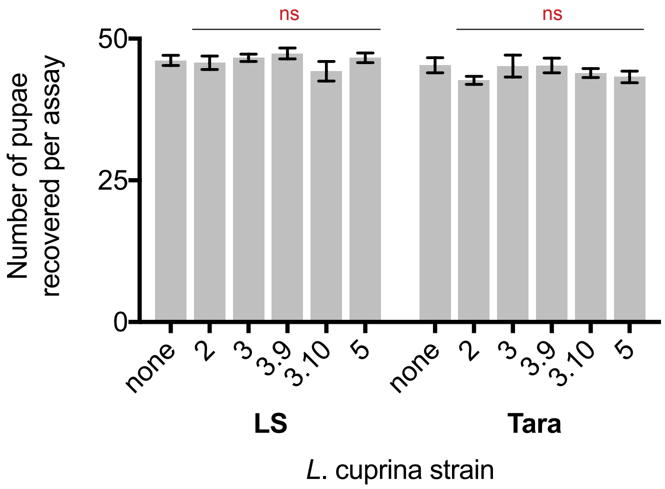

**Supplementary Figure 12. Boronic acid compounds had no effect on *Lucilia cuprina* pupation in the absence of the organophosphates diazinon or malathion.** The number of pupae recovered in the absence of boronic acids is compared to the number recovered in the presence of boronic acids at 1 mg per assay for both LS and Tara *L. cuprina* strains. Data is mean ± SEM for three replicate experiments each starting with 50 larvae (ns = not significant; one-way ANOVA to followed by Dunnett's multiple comparison test).

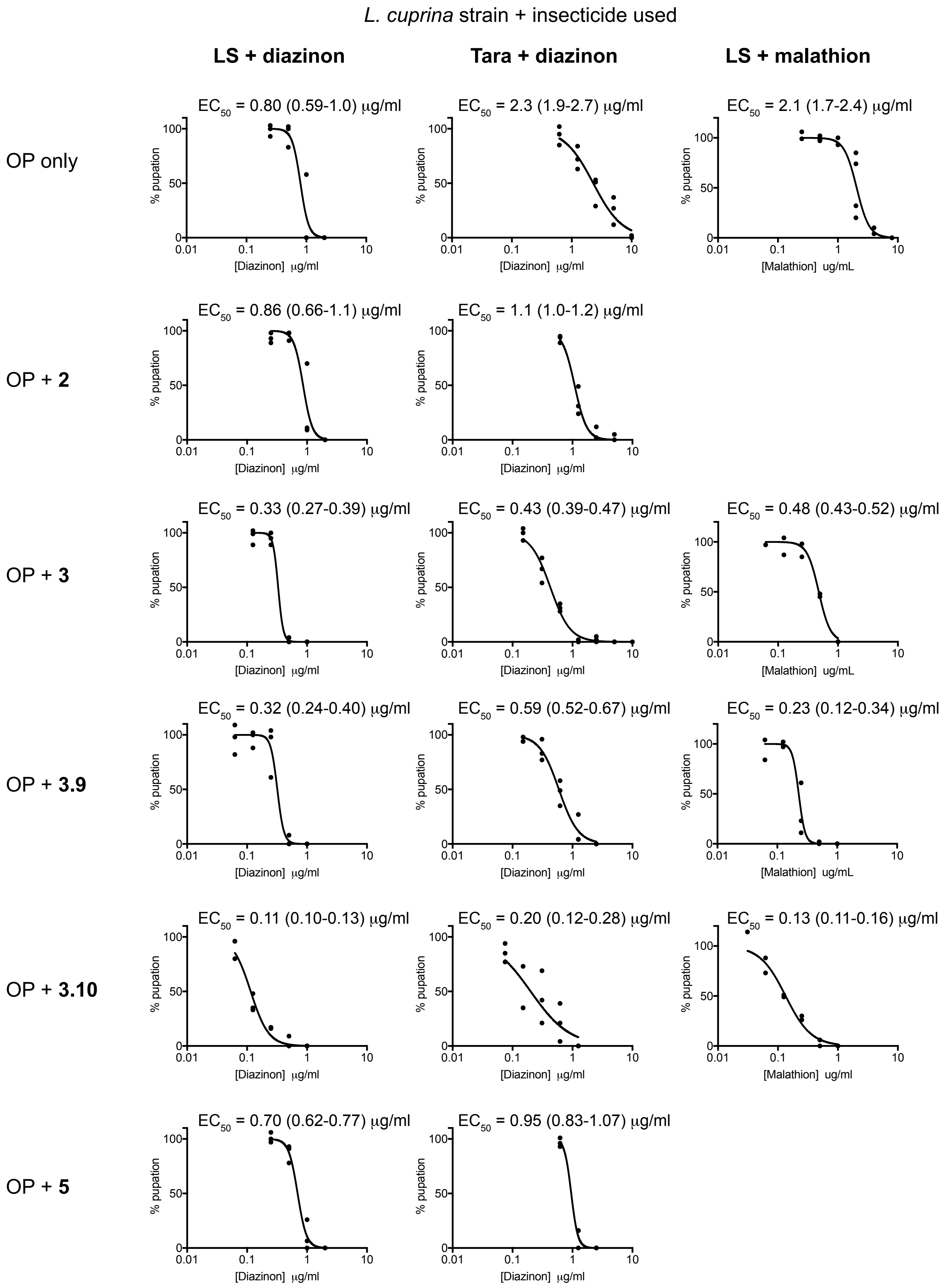

**Supplementary Figure 13. Dose-response curves for *Lucilia cuprina* bioassay.** Treatment consisted of organophosphate insecticide only (diazinon / malathion) or insecticide supplemented with boronic acid compound at a set concentration of 1 mg/ml. The concentration of organophosphate insecticide required to reduce the pupation rate by 50% ($EC_{50}$) was calculated by fitting a sigmoidal dose-response curve to plots of percentage pupation. Each of the data points represents the percentage pupation for 50 initial blowfly larvae with the bioassay repeated three times (diazinon), or two times (malathion), for each concentration of insecticide. The $EC_{50}$ values are presented ± 95% confidence interval.

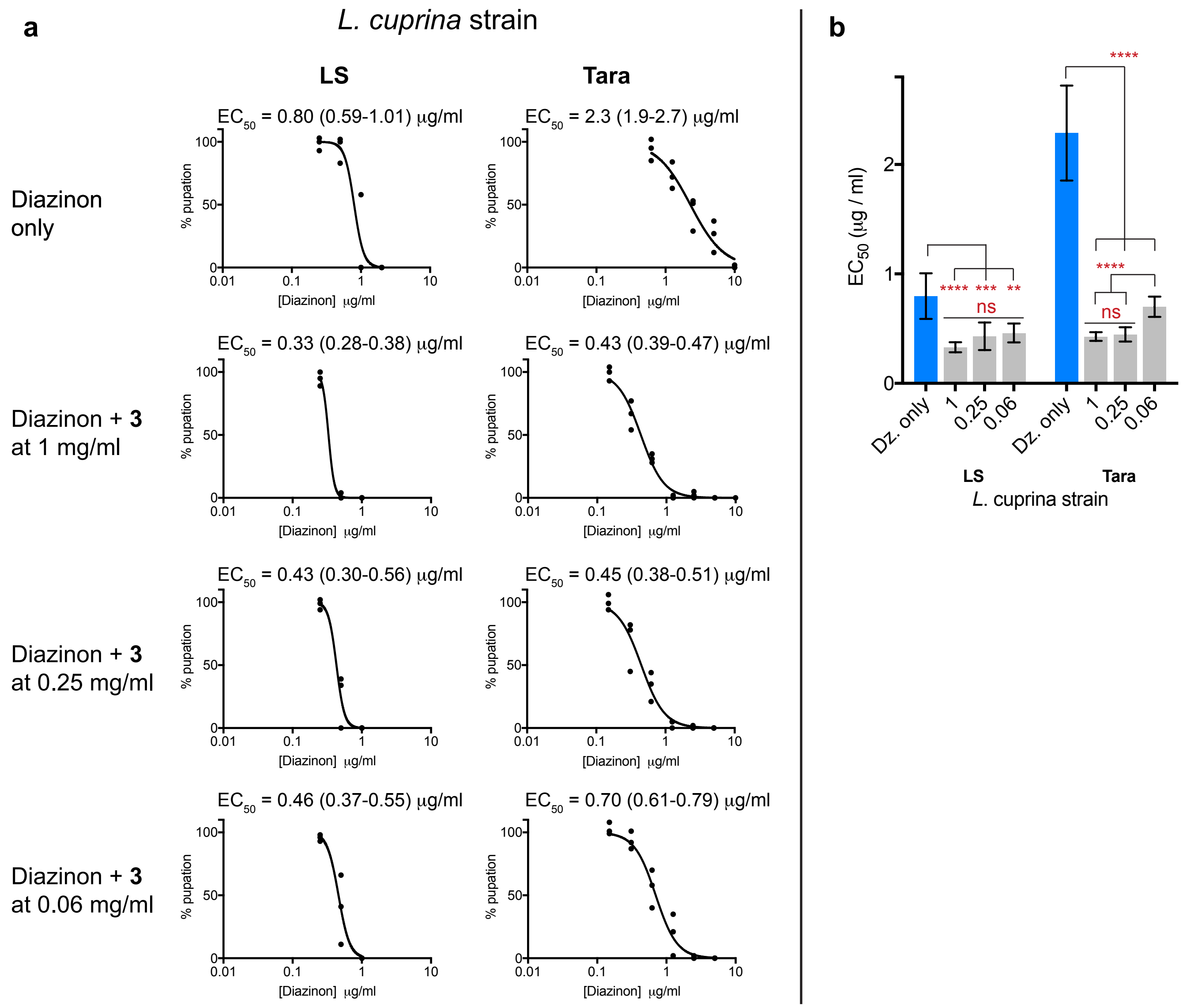

**Supplementary Figure 14. Compound 3 synergizes diazinon against *L. cuprina* at decreased concentrations. (a)** Dose-response curves for *L. cuprina* bioassays at reduced compound concentration. Treatment consisted of diazinon only, or diazinon supplemented with boronic acid compound at a set concentration of 1, 0.25 or 0.06 mg/ml. The concentration of diazinon required to reduce the pupation rate by 50% ($EC_{50}$) was calculated by fitting a sigmoidal dose-response curve to plots of percentage pupation. Each of the data points represents the percentage pupation for 50 initial blowfly larvae with the bioassay repeated three times for each concentration of diazinon. The $EC_{50}$ values are quoted as the best-fit value ± 95% confidence interval. **(b)** There was no significant difference between diazinon $EC_{50}$ values when the concentration of compound **3** was reduced to 0.25 and 0.06 mg/ml against the LS blowfly strain, however there was an increase in $EC_{50}$ against the Tara strain when the concentration compound **3** was reduced to 0.06 mg/ml. $EC_{50}$ values are presented ± 95% confidence interval (**$P<0.01$, ***$P<0.001$, ns = not significant; one-way ANOVA to followed by Dunnett's multiple comparison test).

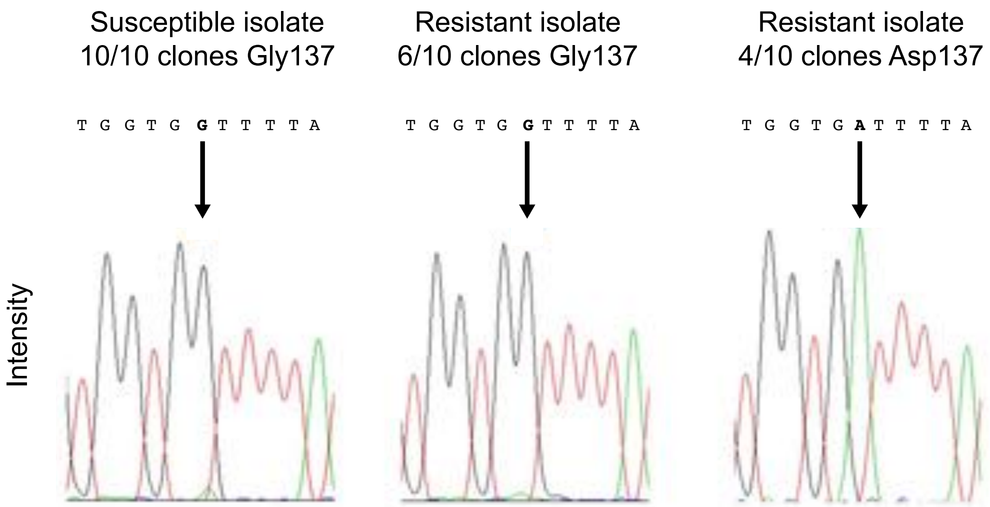

**Supplementary Figure 15. Sequencing of the *Lc*αE7 gene in susceptible (LS) and resistant (Tara) isolates.** The susceptible *Lucilia cuprina* stain only carries the wild type αE7 gene, while the resistant strain contains both wild type αE7 (Gly137) and the Gly137Asp variant. Chromatograms and corresponding nucleotides are shown for the relevant region of the αE7 gene.

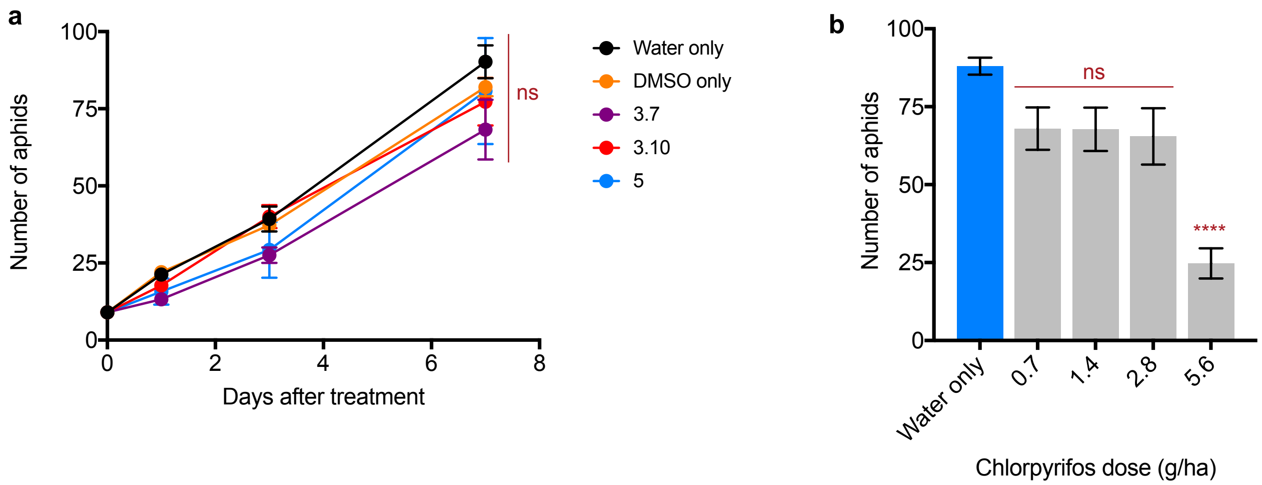

**Supplementary Figure 16. Boronic acid compounds synergize chlorpyrifos against the peach-potato aphid *Myzsus persicae*. (a)** The boronic acid compounds by themselves had no significant effect on the number of aphids compared to a DMSO only control. Data is mean ± SEM for four repeat experiments (ns = not significant; one-way ANOVA followed by Dunnett's multiple comparison test). **(b)** The effect of chlorpyrifos at four doses seven days after treatment. Only treatment with 5.6 g/ha gave a significant decrease in the number of aphids compared to a water only control ( ****P<0.0001, ns = not significant; one-way ANOVA followed by Dunnett's multiple comparison test).

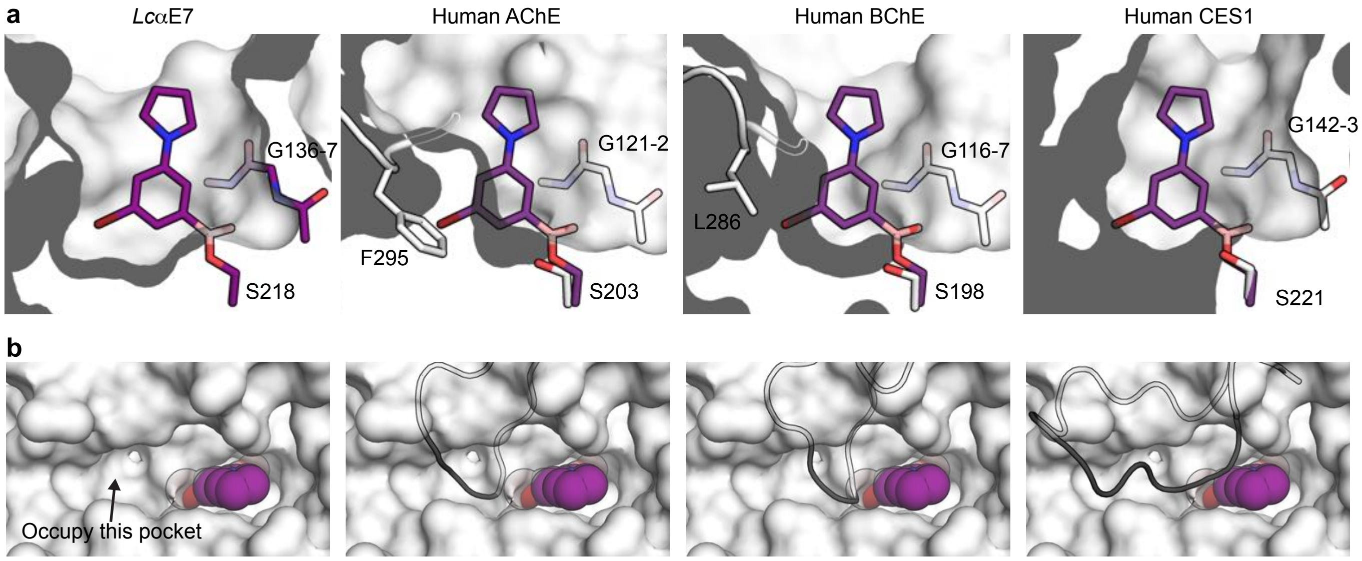

**Supplementary Figure 17. Structural basis for selectivity against human carboxylesterases. (a)** Sliced-surface view of compound **3** overlaid with human acetylcholinesterase (AChE, PDB code 4EY4[11]), human butyrylcholinesterase (BChE, 1P0I[12]) and human carboxylesterase 1 (CES1, PDB code 4AB1[13]). High selectivity against AChE is conferred by a steric clash between Phe295 and the bromo substituent of compound **3**. The enlarged active site of BChE (via replacement of Phe295 with an isoleucine, and a shift in the loop which positions this residue) explains the reduced selectivity of compound **3** against BChE. CES1 has an open active site which allows it to accommodate compound **3**, resulting in low selectivity. **(b)** Surface view of the *Lc*αE7 active site entrance showing a nearby hydrophobic pocket, which if occupied, could confer increased selectivity because of the steric clash with a loop which exists in AChE, BChE and CES1 (black cartoon). Compound **3** is shown as purple spheres.

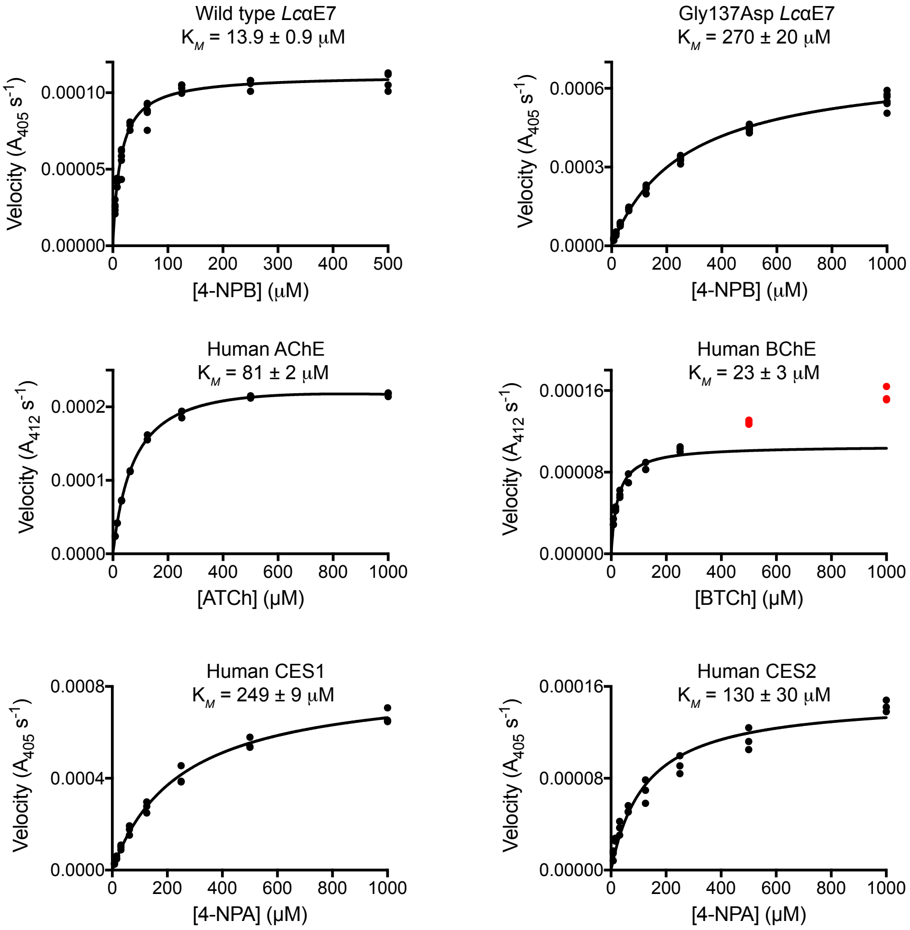

**Supplementary Figure 18. Kinetic parameters for activity assays.** Wild type and Gly137Asp *Lc*αE7 were assayed with 4-nitrophenol butyrate (4-NPB), human acetylcholinesterase (AChE) was assayed with acetylthiocholine (ATCh), human butyrylcholinesterase (BChE) was assayed with butyrylthiocholine (BTCh) and human carboxylesterase 1 and 2 (CES1 and CES2) were both assayed with 4-nitrophenyl acetate (4-NPA). Parameters were determined by fitting the Michaelis-Menten equation to plots of enzyme velocity at eight substrate concentrations using non-linear regression. BChE shows activation at high substrate concentrations[14], hence the velocity measurements at 500 and 1000 μM (red dots) were excluded. At least three repeat measurements of enzyme activity were performed for each concentration of substrate. The Michaelis constant ($K_M$) is presented ± SE.

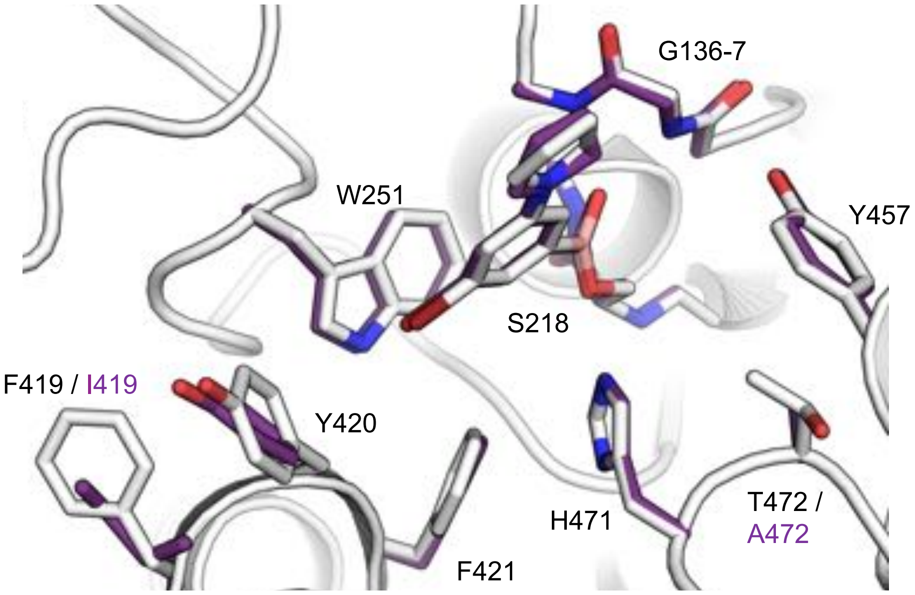

**Supplementary Figure 19. Internal stabilizing mutations present in *Lc*αE7-4a have no effect on boronic acid binding.** The co-crystal structure of compound **3** with *Lc*αE7-4a[15] (white sticks, PDB code 5TYN) is aligned with the co-crystal structure of compound **3** with Asp83Ala + Lys530Glu *Lc*αE7 (purple sticks, PDB code 5TYM). The only structural difference is a 30° rotation in the $\chi_2$ dihedral angle of Tyr420 as a result of the Ile419Phe mutation. For clarity, only the backbone of *Lc*αE7-4a is shown (white cartoon).

**Supplementary Table 1. Data collection and refinement statistics.**

| PDB code (compound) | 5TYP (**1**) | 5TYO (**2**) | 5TYN (**3**) | 5TYL (**4**) | 5TYK (**5**) | 5TYM (**3**) | 5TYJ (**3.10**) |
|---|---|---|---|---|---|---|---|
| **Data collection** | | | | | | | |
| Space group | $C222_1$ | $C222_1$ | $C222_1$ | $C222_1$ | $C222_1$ | $C222_1$ | $C222_1$ |
| Cell dimensions | | | | | | | |
| *a*, *b*, *c* (Å) | 48.46, 101.8, 224.7 | 49.08, 101.2, 222.8 | 49.71, 101.07, 222.6 | 48.77, 101.4, 223.5 | 47.80, 100.8, 221.8 | 49.15, 100.8, 224.0 | 48.58, 102.5, 224.4 |
| α, β, γ (°) | 90, 90, 90 | 90, 90, 90 | 90, 90, 90 | 90, 90, 90 | 90, 90, 90 | 90, 90, 90 | 90, 90, 90 |
| Resolution (Å) | 1.88 (1.92-1.88)* | 1.57 (1.60-1.57) | 1.53 (1.56-1.53) | 1.75 (1.78-1.75) | 1.65 (1.68-1.65) | 1.84 (1.88-1.84) | 1.75 (1.78-1.75) |
| $R_{merge}$ | 0.16 (2.5) | 0.089 (2.7) | 0.16 (1.6) | 0.14 (2.3) | 0.13 (2.2) | 0.14 (2.48) | 0.12 (3.08) |
| $I / \sigma I$ | 11.9 (1.1) | 21.6 (1.0) | 6.9 (1.0) | 10.3 (0.8) | 14.4 (1.3) | 12.4 (1.1) | 13.3 (0.9) |
| Completeness (%) | 99.9 (98.3) | 97.0 (73.8) | 99.5 (97.8) | 100 (99.9) | 100 (99.3) | 99.9 (99.9) | 100 (100) |
| Redundancy | 13.4 (10.9) | 14.3 (11.5) | 6.6 (6.9) | 7.1 (6.9) | 14.0 (13.9) | 11.9 (10.9) | 14.4 (14.5) |
| **Refinement** | | | | | | | |
| Resolution (Å) | 1.88 | 1.57 | 1.53 | 1.75 | 1.65 | 1.84 | 1.75 |
| No. reflections | 45621 | 75341 | 84196 | 56360 | 64943 | 48767 | 57007 |
| $R_{work}$ / $R_{free}$ | 0.221 / 0.283 | 0.206 / 0.236 | 0.193 / 0.233 | 0.214 / 0.263 | 0.187 / 0.223 | 0.228 / 0.287 | 0.198 / 0.240 |
| No. atoms | | | | | | | |
| Protein | 4567 | 4571 | 4614 | 4565 | 4572 | 4564 | 4579 |
| Ligand/ion | 11 | 17 | 14 | 13 | 18 | 14 | 17 |
| Water | 129 | 262 | 421 | 194 | 266 | 117 | 205 |
| *B*-factors | | | | | | | |
| Protein | 40.3 | 27.4 | 20.8 | 30.8 | 25.6 | 35.9 | 39.8 |
| Ligand/ion | 36.4 | 30.0 | 15.9 | 26.9 | 34.0 | 25.9 | 29.8 |
| Water | 33.6 | 29.6 | 27.3 | 29.7 | 28.7 | 30.8 | 37.9 |
| R.m.s. deviations | | | | | | | |
| Bond lengths (Å) | 0.009 | 0.007 | 0.007 | 0.008 | 0.007 | 0.008 | 0.007 |
| Bond angles (°) | 1.2 | 1.0 | 1.0 | 1.1 | 1.1 | 1.1 | 1.0 |

*Values in parentheses are for highest-resolution shell.

**Supplementary Table 2.** ***In vitro*** **inhibition of carboxylesterase enzymes by boronic acid compounds.**

| | ***K*i inhibition constant**[a] | | | | | |
|---|---|---|---|---|---|---|
| **Compound** | Wild type αE7 (**nM**) | G137D αE7 (**nM**) | hAChE (**μM**) | hBuChE (**nM**) | hCES1 (**nM**) | hCES2 (**nM**) |
| PBA | 210 (170-260) | 450 (370-550) | | | | |
| **1** | 12 (9-16) | 49 (42-56) | 81 (65-100) | 260 (220-300) | 0.93 (0.84-1.1) | 120 (91-160) |
| **2** | 2.9 (2.2-3.8) | 29 (27-32) | 63 (50-84) | 310 (260-350) | 1.4 (1.3-1.6) | 9.1 (7.0-12) |
| **3** | 0.25 (0.22-0.28) | 110 (97-130) | 25 (20-31) | 260 (220-320) | 4.1 (3.5-4.9) | 21 (15-28) |
| **4** | 7.2 (6.0-8.7) | 190 (160-230) | 190 (130-310) | 56 (48-66) | 1.8 (1.6-2.1) | 1100 (900-1500) |
| **5** | 11 (9-13) | 170 (140-190) | 26 (21-38) | 370 (300-450) | 3.4 (3.0-3.9) | 59 (45-77) |
| **3.1** | 0.70 (0.62-0.79) | 430 (380-500) | | | | |
| **3.2** | 0.35 (0.31-0.40) | 210 (180-230) | | | | |
| **3.3** | 0.47 (0.36-0.59) | 150 (130-170) | | | | |
| **3.4** | 3.8 (2.9-4.9) | 1000 (900-1200) | | | | |
| **3.5** | 3.4 (2.6-4.5) | 440 (350-550) | | | | |
| **3.6** | 6.3 (4.5-8.8) | 71 (60-85) | | | | |
| **3.7** | 0.30 (0.26-0.35) | 76 (69-85) | | | | |
| **3.8** | 2.0 (1.7-2.4) | 110 (100-120) | | | | |
| **3.9** | 0.44 (0.39-0.50) | 25 (22-27) | 14 (10-19) | 27 (25-30) | 1.1 (1.0-1.2) | 2.2 (1.7-2.8) |
| **3.10** | 0.44 (0.34-0.56) | 18 (16-20) | 9.3 (11-7.7) | 94 (80-110) | 1.1 (1.0-1.3) | 2.1 (1.6-2.8) |
| **3.11** | 5.8 (4.1-8.0) | 1000 (800-1200) | | | | |
| **3.12** | 12 (9-16) | 1000 (900-1100) | | | | |

[a] Inhibition constants are presented with 95% confidence intervals in parentheses. See **Supplementary Fig. 2, 5**, **7** and **8** for dose-response curves.

**Supplementary Table 3. Synergists are selective over human proteases**

| Protease | 3 [a] | 3.9 [a] | 3.10 [a] |
|---|---|---|---|
| **DPP4** | 6 | 2 | -3 |
| **DPP8** | 42 | 53 | 37 |
| **DPP9** | 45 | 75 | 71 |
| **Factor VII** | 15 | 26 | 53 |
| **Factor-Xa** | 53 | 26 | 35 |
| **Furin** | 1 | -2 | -5 |
| **GranzymeA** | 12 | 23 | 12 |
| **GranzymeB** | 3 | 17 | 11 |
| **GranzymeK** | 5 | 14 | 3 |
| **HTRA2** | 23 | 28 | 2 |
| **Kallikrein11** | -71 | -148 | -43 |
| **Kallikrein13** | 3 | -1 | 0 |
| **Matriptase** | -1 | 5 | 4 |
| **Plasma-Kallikrein** | 6 | 41 | 64 |
| **Plasmin** | 13 | 16 | -2 |
| **Prolyl Oligopeptidase** | 61 | 48 | 30 |
| **PSMB5** | 3 | 34 | 2 |
| **PSMB6** | 16 | 54 | 27 |
| **PSMB7** | 39 | 70 | 48 |
| **PSMB8** | 19 | 52 | 49 |
| **PSMB9** | 12 | 50 | 15 |
| **PSMB10** | 43 | 78 | 45 |
| **Spinesin** | 26 | 13 | 34 |
| **Thrombin** | 32 | 54 | 58 |
| **tPA** | 26 | 40 | 19 |
| **uPA** | 67 | 40 | 32 |

[a] Average % inhibition of protease at 100 μM of the indicated boronic acid compound (n=2). Inhibition >50% is marked in red.

**Supplementary Table 4. Protease selectivity panel**

| Target | Vendor | Lot# | [Enzyme], nM | Substrate | Incubation Time (hr) |
|---|---|---|---|---|---|
| DPP4 | ENZO | 2021206 | 0.369 | RPAGARK | 1.5 |
| DPP8 | ENZO | 2281220 | 0.5 | RPAGARK | 1.5 |
| DPP9 | ENZO | 7221432 | 0.3125 | RPAGARK | 1.5 |
| Factor VII[a] | R&D Systems | NJX0716031 | 38.6 | BOC-VPA-AMC | 0.3 |
| Factor Xa | Calbiochem | B76791 | 20 | VMIAALPRTMFIQRR | 5 |
| Furin | R&D Systems | INK025061 | 16 | LRRVKRSLDDA | 6 |
| Granzyme A | ENZO | 11061411 | 30 | PRTLTAKK | 5 |
| Granzyme B | ENZO | 9161556 | 0.493 | IEPDSGGKRK | 5 |
| Granzyme K | ENZO | L27095 | 40 | RPAGARK | 5 |
| HTRA2 | R&D Systems | HVL1015111 | 76.1 | Casein Fluoroscein | 3 |
| Kallikrein 11[b] | R&D Systems | MZV0116081 | 0.032 | BOC-VPA-AMC | 3 |
| Kallikrein 13[c] | R&D Systems | NXS0117021 | 8.9 | BOC-VPA-AMC | 0.2 |
| Matriptase | R&D Systems | PZZ0916101 | 0.3134 | BOC-QAR-AMC | 0.5 |
| Plasma-Kallikrein | R&D Systems | NVH0111081 | 1.5 | ARDIYAASFFRK | 1 |
| Plasmin[d] | R&D Systems | MQB0411091 | 2 | KHPFHLVIHTKR | 2 |
| Prolyl Oligopeptidase | R&D Systems | QBQ0212011 | 0.1 | VMIAALPRTMFIQRR | 2 |
| PSMB5 | BOSTON-BIOCHEM | 21118515B | 0.8 | TYETFKSIMKKSPF | 1.75 |
| PSMB6 | BOSTON-BIOCHEM | 21118515B | 1 | GLTNIKTEEISEVNLDAEFRKKRR | 3.75 |
| PSMB7 | BOSTON-BIOCHEM | 21118515B | 0.24 | GRSRSRSRSR | 2 |
| PSMB8 | BOSTON-BIOCHEM | 04720017A | 0.8 | TYETFKSIMKKSPF | 4.5 |
| PSMB9 | BOSTON-BIOCHEM | 04720017A | 1 | GLTNIKTEEISEVNLDAEFRKKRR | 3.75 |
| PSMB10 | BOSTON-BIOCHEM | 04720017A | 2 | GRSRSRSRSR | 2 |
| Spinesin[b] | R&D Systems | NOS031701A | 8.721 | BOC-QAR-AMC | 0.5 |
| Thrombin | R&D Systems | HWO0413121 | 1 | PRTLTAKK | 1.5 |
| tPA | R&D Systems | DATN0212041 | 9 | VMIAALPRTMFIQRR | 6 |
| uPA | R&D Systems | HKY0112041 | 6.5 | ADFVRAARR | 3 |

[a] Cascade assay where enzyme was first activated by Thermolysin and then Factor III.
[b] Cascade assay where enzyme was activated by Thermolysin.
[c] Cascade assay where enzyme was activated by Lysyl Endopeptidase.
[d] Cascade assay where enzyme was activated by 0.04x of uPA.

**Supplementary Table 5. Effects of boronic acid inhibitors on sensitivity of blowfly larvae to diazinon and malathion.**

| Blowfly strain | Drug treatment[a] | Organophosphate dose response | | | |
|---|---|---|---|---|---|
| | | $EC_{50}$[b] (µg/assay) | 95% CI | SR[c] | RR[d] |
| Laboratory (LS) | Dz. alone | 0.80 | 0.59 – 1.01 | - | - |
| | Dz. plus **2** | 0.86 | 0.66 – 1.06 | 0.9 | - |
| | Dz. plus **3** | 0.33 * | 0.27 – 0.39 | 2.4 | - |
| | Dz. plus **3** (0.25 mg/ml) | 0.43 * | 0.30 – 0.56 | 1.9 | - |
| | Dz. plus **3** (0.05 mg/ml) | 0.46 * | 0.37 – 0.55 | 1.7 | - |
| | Dz. plus **3.9** | 0.32 * | 0.25 – 0.40 | 2.5 | - |
| | Dz. plus **3.10** | 0.11 * | 0.10 – 0.13 | 7.3 | - |
| | Dz. plus **5** | 0.70 | 0.62 – 0.77 | 1.1 | - |
| Field (Tara) | Dz. alone | 2.29 | 1.85 – 2.72 | - | 2.86 |
| | Dz. plus **2** | 1.09 * | 0.99 – 1.18 | 2.1 | 1.36 |
| | Dz. plus **3** | 0.43 * | 0.39 – 0.47 | 5.3 | 0.54 |
| | Dz. plus **3** (0.25 mg/ml) | 0.45 * | 0.38 – 0.51 | 5.1 | 0.56 |
| | Dz. plus **3** (0.05 mg/ml) | 0.70 * | 0.61 – 0.79 | 3.3 | 0.88 |
| | Dz. plus **3.9** | 0.59 * | 0.52 – 0.67 | 3.9 | 0.74 |
| | Dz. plus **3.10** | 0.20 * | 0.12 – 0.28 | 11.5 | 0.25 |
| | Dz. plus **5** | 0.95 * | 0.83 – 1.08 | 2.4 | 1.19 |
| Laboratory (LS) | Mal. alone | 2.06 | 1.73 – 2.39 | - | - |
| | Mal. plus **3** | 0.48 * | 0.43 – 0.52 | 4.3 | - |
| | Mal. plus **3.9** | 0.23 * | 0.12 – 0.34 | 9.0 | - |
| | Mal. plus **3.10** | 0.13 * | 0.11 – 0.16 | 15.8 | - |

[a] Diazinon (Dz.) and malathion (Mal.) examined at a range of concentrations in the presence or absence of boronic acid at constant concentration of 1 mg / ml (except where indicated)
[b] * indicates that, within a blowfly strain, $EC_{50}$ for Dz./Mal. plus boronic acid is significantly different from $EC_{50}$ for Dz./Mal. alone (*P<0.0001; one-way ANOVA to followed by Dunnett's multiple comparison test)
[c] SR = synergism ratio = $EC_{50}$ for Dz./Mal. alone / $EC_{50}$ for Dz./Mal. plus boronic acid, within a blowfly strain
[d] RR = resistance ratio = $EC_{50}$ for Dz. alone or Dz. plus boronic acid with the field strain / $EC_{50}$ for Dz. alone for the laboratory strain
